# Unraveling the therapeutic mechanism of deep-brain stimulation

**DOI:** 10.1101/2022.12.23.521799

**Authors:** Bastijn J.G. van den Boom, Alfredo Elhazaz Fernandez, Peter A. Rasmussen, Enny H. van Beest, Aishwarya Parthasarathy, Damiaan Denys, Ingo Willuhn

**Affiliations:** Netherlands Institute for Neuroscience, Royal Netherlands Academy of Arts and Sciences, Amsterdam, the Netherlands; Department of Psychiatry, Amsterdam UMC, University of Amsterdam, Amsterdam, the Netherlands

## Abstract

Deep-brain stimulation (DBS) is an effective treatment for patients suffering from otherwise therapy-resistant psychiatric disorders, including obsessive-compulsive disorder. Modulation of cortico-striatal circuits has been suggested as a mechanism of action. To gain mechanistic insight, we monitored neuronal activity in cortico-striatal regions in a mouse model for compulsive behavior, while systematically varying clinically-relevant parameters of internal-capsule DBS. DBS showed dose-dependent effects on both brain and behavior: An increasing, yet balanced, number of excited and inhibited neurons was recruited, scattered throughout cortico-striatal regions, while compulsive grooming decreased. Such neuronal recruitment did not alter basic brain function such as resting-state activity, and only occurred in awake animals, indicating a dependency on network activity. In addition to these widespread effects, we observed specific involvement of the medial orbitofrontal cortex in therapeutic outcomes, which was corroborated by optogenetic stimulation. Together, our findings provide mechanistic insight into how DBS exerts its therapeutic effects on compulsive behaviors.

## INTRODUCTION

Electrical deep-brain stimulation (DBS) is used to treat a growing list of neurological and psychiatric disorders^1^. In psychiatry, the fiber bundle most commonly stimulated is the anterior limb of the internal capsule (IC)^2^, often to treat obsessive-compulsive disorder (OCD), a disorder characterized by unwanted thoughts (obsessions) and repetitive behaviors (compulsions)^3,4^. OCD patients that are resistant to conventional therapy benefit substantially from DBS^5–8^. However, the neurobiological mechanism behind DBS remains poorly understood. This poor understanding is reflected in limited therapeutic effect size and long periods of DBS-parameter optimization through trial and error^9^.

An influential theory likens the similar clinical efficacy of DBS to that of capsulotomy^10^, postulating that DBS inhibits surrounding neural tissue and thereby creates a reversible lesion locally^11^. In contrast, several studies point to dysfunctional activity in distal cortico-striatal circuits in OCD^12–14^, and DBS is thought to correct such dysfunction via recruitment of cortical regions^2,15^. Support for this hypothesis stems from patient studies that observed that DBS alters OFC activity^16–18^, modulates frontal theta oscillations^19^, and restores frontostriatal network activity^20^.

To elucidate the mechanism by which DBS acts in cortical and striatal regions, we employed SAPAP3 mutant mice (SAPAP3^-/-^), the best-established model for OCD. These mice exhibit compulsive-like grooming, anxiety-like behavior^21,22^, cognitive deficits^23–26^, and respond well to OCD pharmacotherapy^22^ and DBS^27^. To avoid DBS-induced electrical artifacts, we used calcium imaging to monitor single-cell activity *in vivo*^28^. We applied DBS to the rodent homolog of the human ventral anterior IC, the mouse ventral IC, which carries similar cortical projection fibers^29^. Across experiments, we systematically varied clinically-relevant DBS parameters (current, pulse width, and frequency) in SAPAP3^-/-^ and their wild-type littermates (WT)^9^. We found a DBS dose-dependent reduction in compulsive grooming, accompanied by both brain-wide neuronal dynamics as well as specific responses in the medial orbitofrontal-cortex that were involved in the mechanism by which DBS ameliorates compulsivity.

## RESULTS

### Internal-capsule deep-brain stimulation (IC-DBS) decreases excessive grooming

We examined the effects of DBS parameters on compulsive-like grooming behavior in SAPAP3^-/-^ (*n*=30) and WT (*n*=28) (Supplementary Fig. 1a,b,c). Animals were tested in an open-field apparatus after implanting DBS electrodes into the IC (Fig. 1a,b,c, Supplementary Fig. 1d). All mice were stimulated across three different experiments (on different days) to examine the effects of different intensities (no DBS, low-, medium-, and high-intensity DBS) of clinically relevant DBS parameters (current, pulse width, and frequency) on brain and behavior (Fig. 1d,e). Consistent across experiments, SAPAP3^-/-^ spent approximately 20% of the open-field session grooming at baseline (no DBS), whereas WT groomed only for 5% (Fig. 1f,h). For each of the three DBS-parameter experiments, SAPAP3^-/-^ grooming was not reduced during the lowest DBS-intensity condition (pre-DBS baseline vs DBS; green, 100 μA: t(26)=1.78, p=0.087; 40 μs: t(27)=0.52, p=0.608; 60 Hz: t(25)=1.31, p=0.203;). However, both medium- (yellow) and high-intensity (red) DBS conditions showed immediate and robust reductions in excessive grooming (pre-DBS baseline vs DBS; 200 μA: t(26)=2.34, p=0.040; 300 μA: t(26)=4.44, p=0.003; 80 μs: t(27)=2.35, p=0.039; 160 μs: t(27)=4.18, p=0.005; 120 Hz: t(25)=2.27, p=0.048; 180 Hz: t(25)=2.94, p=0.014) (Fig. 1f). Excessive grooming rapidly reinstated upon DBS offset. Both current and pulse-width experiments exhibited a dose-dependent reduction in grooming (current: F(3,78)=7.82, p<0.001; pulse width: F(3,81)=7.02, p<0.001), whereas increasing frequency beyond 120 Hz did not improve efficacy further (frequency: F(3,75)=1.97, p=0.126) (Fig. 1g). Importantly, DBS did not alter WT grooming (Fig. 1h) or general locomotion (Supplementary Fig. 1f,g). DBS-electrode location (two-dimensional anterior-posterior and dorsal-ventral position) did not correlate with grooming reduction, indicating that precise electrode positioning within the IC did not change effectiveness of DBS (Supplementary Fig. 1e). Exploring novel DBS parameters (low frequency and “cyclic” stimulation) did not result in grooming reduction, in line with recent patient findings^30^ (Supplementary Fig. 1h,i). Taken together, IC-DBS reduced excessive grooming in SAPAP3^-/-^ and, similar to clinical practice, the effectiveness of DBS was improved by adapting current and pulse width, but not frequency^9^.

**Fig. 1.**
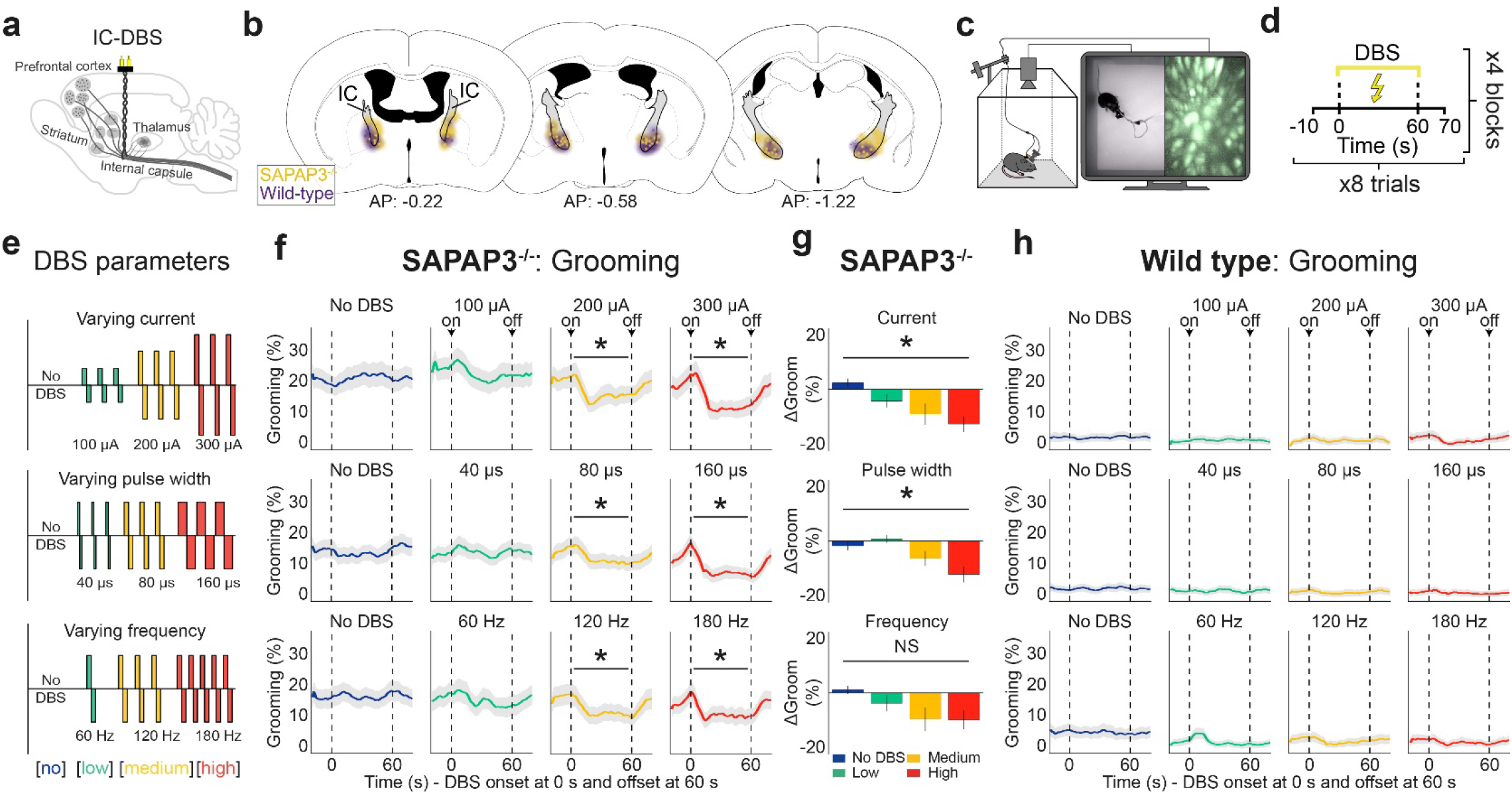
Internal-capsule deep-brain stimulation (IC-DBS) dose-dependently reduces excessive grooming. **a**, Schematic depicting DBS electrodes in the IC, a white-matter bundle that carries corticofugal fibers. **b**, Histological^79^ verification of IC-DBS electrode tips in the IC (gray) of SAPAP3^-/-^ (yellow, *n*=30) and WT (purple, *n*=28). Halo represents the modeled sphere of current spread around the DBS electrode tips. **c**, Mice were subjected to DBS, calcium imaging, and behavioral recordings after being placed in an open field. **d**, DBS was switched ON for 60 s per trial, eight trials per block, and four blocks per session. **e**, Across three sessions, animals were stimulated with varying current (top), pulse width (middle), or frequency (bottom). **f**, DBS reduced compulsive-like grooming during current (top), pulse width (middle), and frequency (bottom) dose-response experiments. **g**, A significant dose-dependent reduction of grooming was observed during current and pulse-width, but not frequency, manipulations. **h**, DBS did not reduce grooming in WT mice. \**p*<0.05, NS=not significant.

### IC-DBS modulates the entire dorsal cortex, with an emphasis on the frontal cortex

DBS is thought to recruit cortical regions^15^. To assess widespread cortical effects of IC-DBS, we employed wide-field calcium imaging across the entire dorsal cortex in *Thy1*-GCaMP6f mice (*n*=5)^31^. The skull was made transparent for calcium imaging^32^ and IC-DBS electrodes implanted, targeted at the ipsilateral hemisphere (Fig. 2a, Supplementary Fig. 2a). After motion correction (Supplementary Fig. 2b,c), the Allen-brain atlas was used to map neuronal activity onto specific brain regions^33^ (Fig. 2b). Similar to the dose-dependent reduction in grooming during varying current and pulse-width experiments (Fig. 1g), we found dose-dependent suppression of the entire dorsal cortex (current: F(3,12)=4.38, p=0.027; pulse width: F(3,12)=7.20, p=0.005), which was absent in the frequency experiment (frequency: F(3,12)=0.31, p=0.845) (Fig. 2c). Analyzing brain regions independently, we found increased activity in all recorded regions immediately upon DBS onset (Fig. 2d). However, within a few seconds activity diminished and sustained suppression of activity was found in a subset of regions. In the frontal cortex (FC), we found suppression across current (F(3,12)=8.77, p=0.005) and pulse-width (F(3,12)=9.64, p=0.005) experiments, and in the somatosensory cortex (SS) only during pulse-width manipulations (F(3,12)=6.41, p=0.023). We found no significant suppression in retrosplenial cortex (RSP) nor visual cortex (VIS), suggesting a rostral-caudal gradient of suppression (Fig. 2d). Direct comparison between regions revealed prominent suppression in FC during current (F(3,12)=4.20, p=0.023; post-hoc: FC vs RSP p=0.043, FC vs VIS p=0.034) and pulse-width (F(3,12)=3.51, p=0.040; post-hoc: FC vs VIS p=0.040) experiments, but not frequency (F(3,12)=2.95, p=0.065) (Fig. 2e). These data indicate that IC-DBS sustainedly suppresses the entire dorsal cortex with a rostral-to-caudal gradient.

**Fig. 2.**
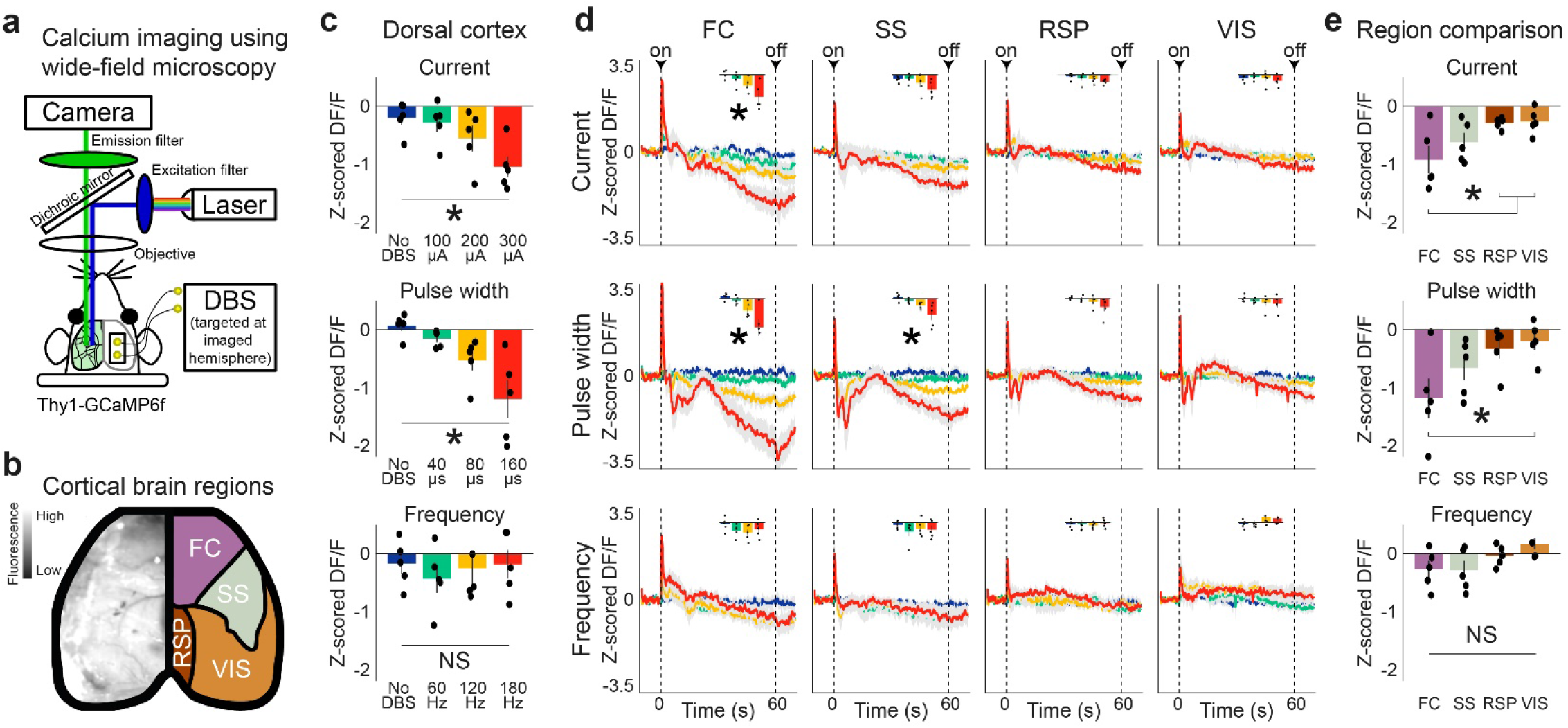
Wide-field calcium imaging reveals IC-DBS modulation of the entire dorsal cortex, with an emphasis on the FC. **a**, Schematic of wide-field fluorescence microscopy setup. DBS electrodes were implanted, inserted at an angle, contralaterally to the imaged hemisphere, and targeted the ipsilateral IC of *Thy1*-GCaMP6f mice (*n*=5). **b**, A video frame of the dorsal cortex (left) and the cortical regions (FC: frontal cortex; RSP: retrosplenial cortex; SS: somatosensory cortex; VIS: visual cortex) as defined by the Allen-brain atlas (right). **c**, Significant dose-dependent cortex-wide suppression was observed in the varying current and pulse-width experiments, but not in the frequency experiment. **d**, The dose-response manipulations of DBS induced region-specific reduction in FC (current and pulse width) and SS (pulse width), but not RSP nor VIS. **e**, Directly comparing sustained suppression at high-intensity DBS across dorsal-cortical regions revealed significant effects during current and pulse-width, but not frequency, experiments. Post-hoc analyses revealed differences between FC and RPS, and FC and VIS during current manipulations. During pulse-width manipulations, FC differed from VIS. \**p*<0.05, NS=not significant.

### Single-cell recruitment in cortical and striatal regions by IC-DBS

Wide-field imaging captures a large part of the brain, but lacks single-cell resolution and is limited to superficial cortical layers in head-fixed mice^34^. To overcome these limitations and elucidate DBS effects on grooming in OCD-relevant circuits, we used miniaturized fluorescent microscopes (miniscopes, Fig. 3a)^35^. Consistent with literature, our wide-field data indicated a role for the FC in IC-DBS^20^. Therefore, we used miniscopes to record from pyramidal neurons in prefrontal cortical regions (lateral and medial orbitofrontal cortex (lOFC, mOFC), prelimbic cortex (PL), and premotor cortex (M2)) and medium-spiny neurons in their striatal projection targets (dorsal and ventral striatum (DS, VS)) (Fig. 3b, Supplementary Fig. 3a). During open-field experiments with freely-behaving SAPAP3^-/-^ and WT, we recorded fluorescence in hundreds of neurons per region and found complex dynamics (Fig. 3c, Supplementary Fig. 3b,c). A subset of neurons exhibited immediate, transient effects (either excitatory or inhibitory) upon DBS onset (Fig. 3d, Supplementary Fig. 3d), resembling cortex-wide increases in activity at DBS onset (Fig. 2d). Other neurons showed sustained excitation or inhibition of their activity during DBS (Fig. 3e, Supplementary Fig. 3e), resembling the sustained suppression we found in the FC (Fig. 2d). In all recorded brain regions, DBS recruited single cells (i.e., modulated their activity) dose-dependently by exciting or inhibiting their activity (Factor intensity, lOFC: F(3,40)=5.53, p=0.003; mOFC: F(3,32)=12.72, p<0.001; PL: F(3,32)=11.27, p<0.001; M2: F(3,32)=14.54, p<0.001; DS: F(3,24)=2.57, p=0.077; VS: F(3,32)=8.75, p<0.001) (Fig. 3f, Supplementary Fig. 3g). However, mOFC and DS predominantly recruited neurons by exciting their activity, independent of genotype (Factor direction, mOFC: F(3,32)=14.77, p<0.001; DS: F(3,24)=7.68, p=0.011). Neurons that were recruited by DBS showed consistency in duration (transient or sustained) and direction (excited or inhibited) across different DBS intensities in SAPAP3^-/-^ (Fig. 3g) and WT (Supplementary Fig. 3f) (e.g., a sustainedly excited recruited neuron during one DBS intensity is likely to be recruited as sustainedly excited again during another DBS intensity), suggesting that DBS affects neurons similarly across different stimulation parameters. Since behavioral DBS effects were restricted to periods of stimulation, we hypothesized that the sustained neurons were driving the recorded reduction in grooming in SAPAP3^-/-^. To investigate whether the sustained activation is driven by direct antidromic stimulation or underlying network activity, we imaged mice under both awake and anesthetized (diminished network activity) conditions in the same session^36^. Under anesthesia, we found no sustained neurons (either excited or inhibited), nor transient inhibited neurons (Fig. 3h), demonstrating that sustained recruitment is dependent on network activity in the awake state. Similar to the consistent recruitment of neurons (direction and duration), we found that transient excited neurons recruited under anesthesia were more likely to be recruited as transient excited neurons in awake recordings (bootstrap, p=0.002) (Fig. 3i), suggesting antidromic stimulation of their axons. Together, DBS modulated neurons*’* activity in a dynamic, yet consistent fashion that was dependent on network activity.

**Fig. 3.**
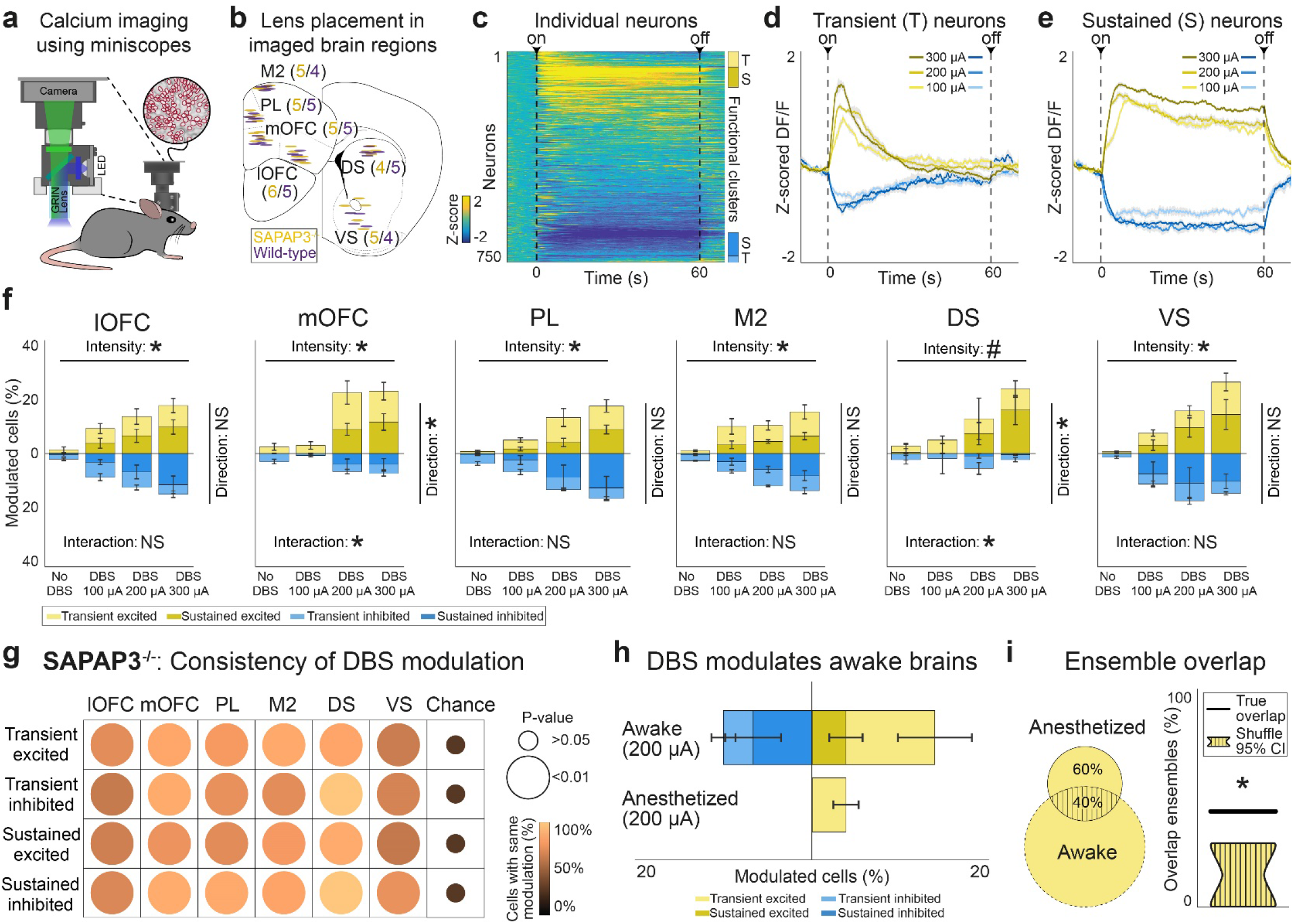
IC-DBS recruits neurons in SAPAP3^-/-^ in cortical and striatal regions via transient/sustained excitation or inhibition. **a**, Calcium imaging in freely-moving mice using miniscopes. **b**, Elliptic shapes represent imaged areas under GRIN lenses in SAPAP3^-/-^ (yellow, *n*=30) and WT (purple, *n*=28)^79^. **c**, Heatmap of all recorded neurons in lOFC (across all SAPAP3^-/-^ mice) during high-intensity DBS, sorted by modulation. Vertical bars on the right depict functional clusters: transient excited (T, light yellow), sustained excited (S, dark yellow), sustained inhibited (S, dark blue), and transient inhibited (T, light blue). **d**, Averaged traces of transiently excited and inhibited neurons during different DBS parameters. **e**, Same as d, but for sustainedly recruited neurons. **f**, Dose-dependent recruitment of excited and inhibited neurons by DBS in SAPAP3^-/-^ were balanced in number in most regions (lOFC, PL, M2, and VS). However, we found an imbalance in the number of recruited excited and inhibited neurons in mOFC and DS. **g**, Stability of direction (excitation or inhibition) and duration (transient or sustained) of single-cell recruitment (i.e., consistency of modulation) was significantly different from chance in all recorded cortical and striatal regions. **h**, Under anesthesia, no sustained neurons (either excited or inhibited), nor transient inhibited neurons, were found. **i**, Overlap of identity of transiently excited neurons found between awake and anesthetized states was significantly different from chance (95% confidence intervals presented). \**p*<0.05, #0.05<*p*<0.1, NS=not significant.

### Unaffected basic brain function during IC-DBS

To examine if DBS alters all aspects of neuronal activity, we quantified proxies of “basic brain function”: we calculated the baseline activity of neurons (a form of resting-state activity)^37^ and examined how anatomical distance between neurons affected the synchrony of their activity (spatiotemporal correlations)^38^. First, we compared averaged regional baseline activity during the “no-DBS” block with baseline activity during the high-intensity DBS block. The cumulative density function of calcium events and the average frequency of calcium events were combined into a cell-activity index measure (Fig. 4a). The cell-activity index during the high-intensity DBS block was not different from the “no-DBS” block in SAPAP3^-/-^ (Fig. 4b), nor WT (Supplementary Fig. 4a), indicating that resting-state activity was not affected by DBS. Next, we tested whether spatiotemporal correlations (i.e., distance between neuron pairs correlated with their activity)^39^ were preserved during DBS (Fig. 4c). Local spatiotemporal correlations between recorded neurons were found in all regions during the “no-DBS” block and were preserved during DBS (Fig. 4d). Since DBS strongly modulated neurons*’* activity (Fig. 3f), we reasoned that preserved local spatiotemporal correlations would likely be achieved by scattered recruitment of neurons (i.e., lack of spatial clustering). To properly assess anatomical organization (Supplementary Fig. 4b), we calculated the anatomical distance of each recruited neuron to its closest recruited neighbor (to avoid averaging out short and long distances between neuron pairs), tested to chance (Fig. 4e), and did not find clustering of neurons (Fig. 4f). Together, these data indicate that, although DBS recruits neurons widespread throughout brain regions, it does not compromise basic brain function.

**Fig. 4.**
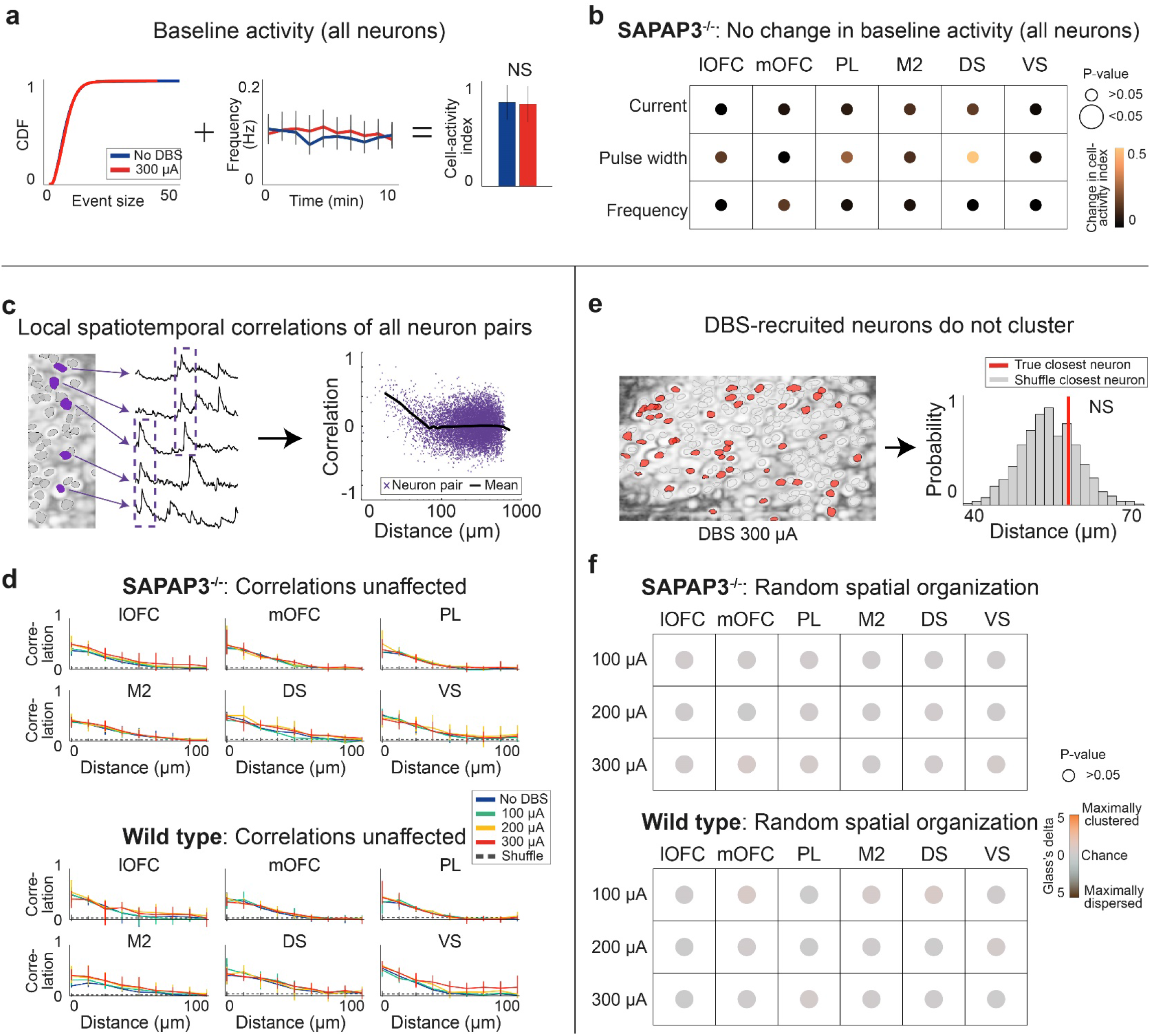
Basic brain function not affected by IC-DBS. **a**, Cumulative distribution function (left) and averaged frequency of firing (middle) during the “no-DBS” block and the high-intensity DBS block were combined into a single neuronal-activity measure: Cell-activity index (right), which did not differ between the two blocks. **b**, High-intensity DBS did not change the cell-activity index compared to the “no-DBS” block in SAPAP3^-/-^. **c**, Representative neurons (purple) show synchronous activity when proximal to one another (left). Temporal activity correlation plotted as a function of inter-cell distance (purple, individual neuron pairs; black, average). Distance was logarithmically scaled for visualization. **d**, For both SAPAP3^-/-^ and WT, local spatiotemporal correlations were conserved during DBS, as DBS did not differ from the “no-DBS” block. **e**, Maximum-intensity projection displaying the location of recruited neurons in red circles (left). Example of averaged distance to closest recruited neuron (red) compared to histogram of expected closest neighbor by chance (bootstrap, gray) (right). **f**, No statistically significant clustering for SAPAP3^-/-^ and WT was found in any of the recorded cortical and striatal regions for any of the DBS parameters. NS=not significant.

### Dose-dependent recruitment of similar neuron populations

Next, we focused our analyses on neurons that were recruited sustainedly throughout the stimulation epoch, because their activity change paralleled the duration of grooming reduction and was dependent on network activity (in the awake state). Sustained neurons (both excited and inhibited) were found throughout the miniscope*’*s field-of-view in each DBS-intensity block (Fig. 5a). In all regions, we found dose-dependent recruitment of sustained neurons that amounted to a maximum of approximately a quarter of all imaged neurons in both SAPAP3^-/-^ (lOFC: F(3,15)=7.93, p=0.002; mOFC: F(3,12)=16.56, p<0.001; PL: F(3,12)=8.93, p=0.002; M2: F(3,12)=31.37, p<0.001; DS: F(3,9)=4.32, p=0.038; VS: F(3,12)=5.27, p=0.015) and WT (lOFC: F(3,12)=31.24, p<0.001; mOFC: F(3,12)=11.5, p<0.001; PL: F(3,12)=6.59, p=0.006; M2: F(3,9)=5.5, p=0.020; DS: F(3,12)=7.77, p=0.004; VS: F(3,9)=13.13, p=0.001) (Fig. 5b). Importantly, when comparing SAPAP3^-/-^ with WT, we found stronger recruitment in mOFC in SAPAP3^-/-^ in each DBS-parameter experiment (current: F(1,16)=4.61, p=0.048; pulse width: F(1,16)=9.96, p=0.018; frequency: F(1,16)=7.26, p=0.032) (Fig. 5c), suggesting that the mOFC in SAPAP3^-/-^ was affected more robustly by DBS than WT and potentially drove the suppression of compulsive-like grooming. Since DBS recruited overlapping transient-neuron populations in awake and anesthetized mice (Fig. 3i), we tested whether DBS would also recruit overlapping sustained neuron populations across DBS-intensity blocks (Fig. 5d). Neurons recruited during medium- and high-intensity overlapped above chance level in both SAPAP3^-/-^ (bootstrap - lOFC: p=0.003; mOFC: p=0.003; PL: p=0.003; M2: p=0.003; DS: p=0.012; VS: p=0.024) and WT (bootstrap - lOFC: p=0.003; mOFC: p=0.002; PL: p=0.003; M2: p=0.005; DS: p=0.003; VS: p=0.003) (Fig. 5e, Supplementary Fig. 5a,b). These data are consistent with our behavioral results (Fig. 1f) - only medium- and high-intensity DBS reduced grooming. Since neurons can be excited or inhibited by DBS modulation, we investigated whether a computed excitation/inhibition (E/I) balance observed at baseline was maintained during DBS (Fig. 5f). In most regions, we found preserved E/I balance, as demonstrated by similar number of neurons recruited as excited and inhibited (Fig. 5g). However, during DBS, mOFC and DS neurons were predominantly excited in SAPAP3^-/-^ (mOFC: F(3,16)=3.46, p=0.041; DS: F(3,12)=6.18, p=0.009) and WT (mOFC: F(3,16)=3.92, p=0.028; DS: F(3,16)=5.33, p=0.010), providing more evidence for involvement of mOFC in DBS effects. Together, these data demonstrate “global” DBS effects (which were independent of genotype and brain region): dose-dependent recruitment of neuron populations that partially overlap at therapeutic intensities, while maintaining E/I balance in most regions. These global effects were accompanied by “regional” DBS effects (which were dependent on brain region): the mOFC recruitment in SAPAP3^-/-^, primarily by means of excitation, suggests that mOFC potentially drives the DBS-induced suppression of grooming in SAPAP3^-/-^.

**Fig. 5.**
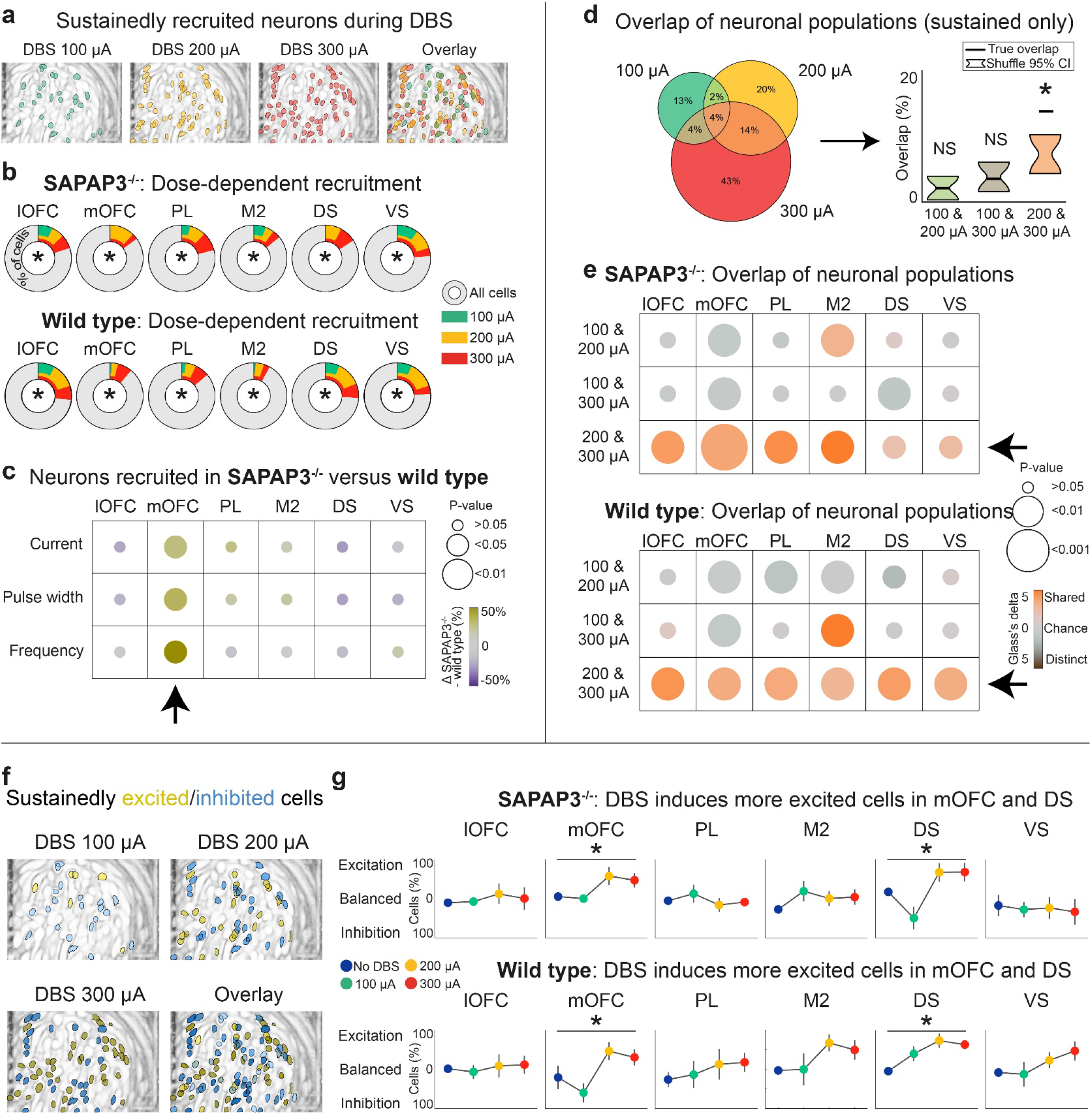
IC-DBS dose-dependently recruits partially overlapping sustained neuron populations, while maintaining excitation/inhibition balance. **a**, Anatomical distribution of imaged neurons revealed by maximum-intensity projection of neurons recruited during 100 μA, 200 μA, 300 μA, and overlay thereof (example animal). **b**, In all recorded regions, we found dose-dependent recruitment of sustained neurons in SAPAP3^-/-^ and WT. **c**, DBS recruited significantly more neurons in mOFC of SAPAP3^-/-^ compared to WT across all stimulation parameters (arrow). **d**, Venn-diagram depicts overlap of neuron populations recruited across different current intensities (example region) (left). True percentage of overlapping neuron populations (black horizontal lines) for the different current blocks compared to chance (95% confidence intervals, bootstrap) indicated recruitment of similar neuron populations during the 200 and 300 μA blocks. **e**, For both SAPAP3^-/-^ and WT, all cortical and striatal regions showed significant overlap in DBS-recruited neuron populations for the 200 and 300 μA blocks (arrows). **f**, Maximum-intensity projection of sustained neurons recruited as excited (yellow) or inhibited (blue) during 100 μA, 200 μA, 300 μA, and overlay thereof (example animal). **g**, For both SAPAP3^-/-^ and WT, we found an increased number of excited neurons in mOFC and DS during DBS. In all other regions, the number of excited and inhibited neurons were balanced during DBS. \**p*<0.05, NS=not significant.

### mOFC in SAPAP3^-/-^ controls compulsive-like grooming

Since DBS reduced grooming specifically, we searched for neurons that were modulated specifically during grooming using Bayesian ANOVAs (Bayes factor>3 to obtain substantial evidence for the presence and absence of effect) (Fig. 6a, Supplementary Fig. 6a). Even using relatively strict Bayesian identification criteria (see Methods), we identified such neurons in all recorded regions (Fig. 6b). For sake of simplicity, we refer to them as grooming-associated neurons. Similarly, we refer to neurons that were not modulated during periods of grooming, locomotion, or behavioral inactivity as not-associated neurons. In the mOFC of SAPAP3^-/-^, we found a trend to dose-dependent recruitment of grooming-associated neurons (mOFC: F(3,16)=3.21, p=0.051) (Fig. 6c, top row). In contrast, in all cortical regions, we found dose-dependent recruitment of not-associated neurons (lOFC: F(3,20)=4.74, p=0.012; mOFC: F(3,16)=3.37, p=0.045; PL: F(3,16)=8.25, p=0.002; M2: F(3,16)=12.86, p<0.001) (Fig. 6c, bottom row), suggesting that DBS does not recruit neurons based on their “cell identity”, which was supported by a lack of overlap between behavior-associated and DBS-recruited neuronal populations (Supplementary Fig. 6b,c,d,e). We hypothesized that DBS reduces grooming by diminishing the number of grooming-associated neurons. This hypothesis is supported by the finding that the number of grooming-associated neurons in mOFC was consistently reduced during DBS across DBS-parameter experiments (mOFC - current: t(4)=4.21, p=0.041; pulse width: t(4)=2.8, p=0.049; frequency: t(4)=3.88, p=0.036) (Fig. 6d), suggesting that the activity of mOFC neurons contributes to grooming frequency. To test whether these neurons causally contribute to reduced grooming, we expressed the excitatory opsin ChETA^40^ in mOFC (*n*=7) to mimic regional excitatory DBS effects (Fig 5g, Fig. 6e, Supplementary Fig. 6f). Photostimulation was delivered for 60 s (to mimic the activity of sustainedly recruited neurons). Indeed, grooming diminished and re-emerged at photostimulation on- and off-set, respectively (473 nm, 5 mW, 10 ms pulse-duration) (Fig. 6f), at 5, 15 and 120 Hz (5Hz: t(6)=3.86, p=0.033; 15Hz: t(6)=4.17, p=0.029; 120: t(6)=3.58, p=0.035). In contrast, 60 s photostimulation at 1 Hz or photostimulation at 15 Hz for 5 s (to mimic the activity of transiently recruited neurons) had no effect on grooming (Fig. 6g). Photostimulation, regardless of protocol, did not affect general locomotion or the relationship between grooming and locomotion (Supplementary Fig. 6g,h). Changes in grooming were abolished when light was prevented from entering the mOFC by blocking the implanted fibers. Importantly, control animals (*n*=5), expressing only the stable fluorophore mCherry in mOFC, and animals expressing ChETA in lOFC (*n*=5) or M2 (*n*=5) did not exhibit reduced grooming. Taken together, mOFC neurons exhibit grooming-related information that is altered by DBS and tightly linked to compulsive-like grooming behavior.

**Fig. 6.**
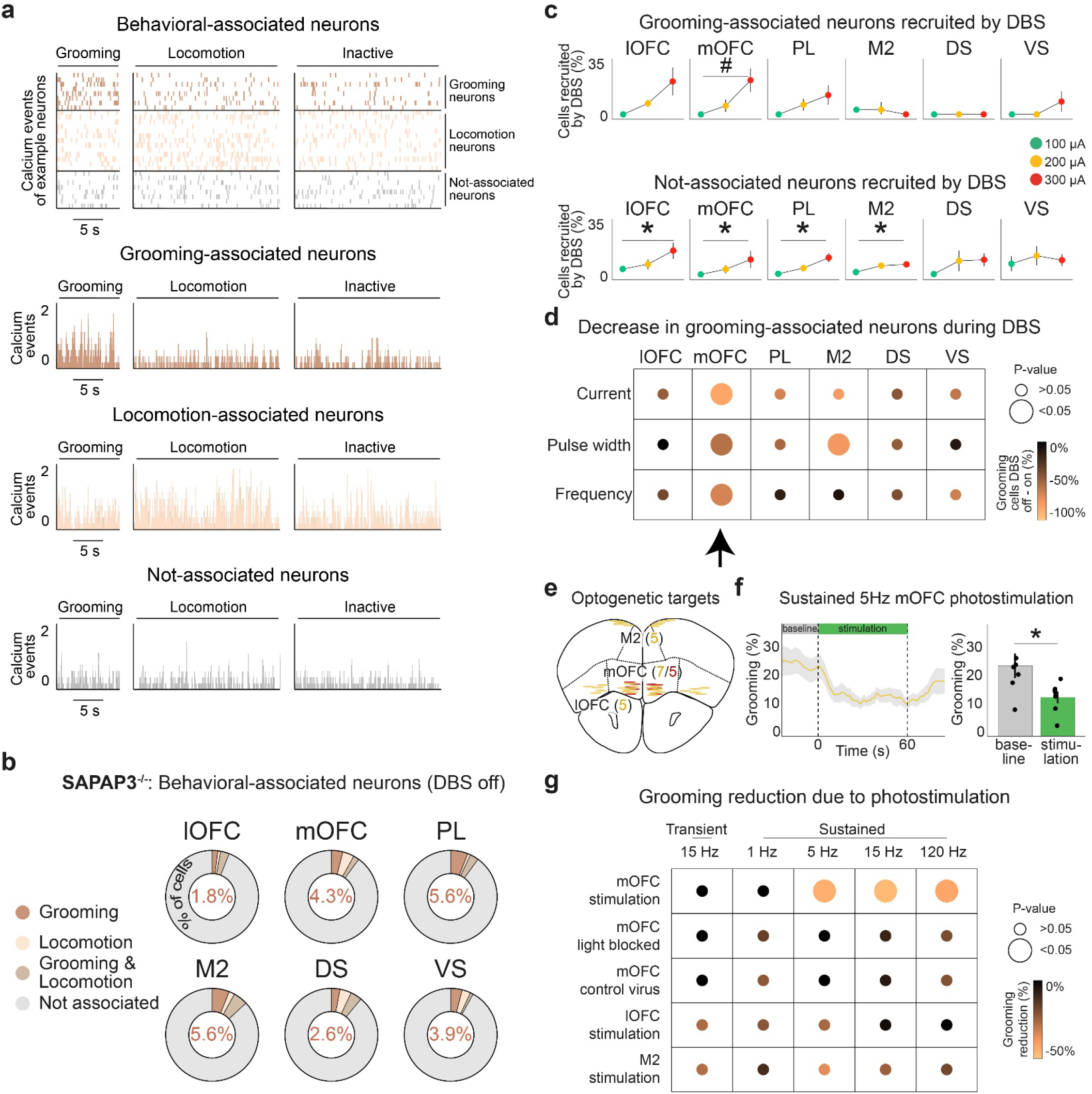
DBS reduced grooming-associated neurons in the mOFC of SAPAP3^-/-^, as validated with optogenetics. **a**, Behavior-associated neurons classified as grooming, locomotion, grooming and locomotion, or not-associated neurons using Bayesian ANOVAs (example animal displayed). Raster plot of the deconvolved calcium events of all behavior-associated neurons across time (top). Histograms show the number of binned calcium events for grooming-associated (top, brown), locomotion-associated (middle, pink), and not-associated (bottom, gray) neurons per behavioral period (i.e., grooming, locomotion, or inactive). **b**, In each recorded cortical and striatal region, we detected grooming-, locomotion-, and grooming&locomotion-associated neurons (percentage of grooming-associated neurons depicted in donut charts). **c**, DBS did not recruit grooming-associated neurons significantly in a dose-dependent manner in any of the recorded regions (top), although mOFC exhibited a trend (p=0.051). DBS did recruit not-associated neurons dose-dependently in all recorded cortical regions (bottom). **d**, DBS reduced the number of grooming-associated neurons in mOFC consistently across stimulation parameters (arrow). **e**, Schematic^79^ of bilateral optogenetic stimulation of the mOFC (*n*=7), lOFC (*n*=5), and M2 (*n*=5) with ChETA (yellow dots), and mOFC (*n*=5) with mCherry (red dots) in SAPAP3^-/-^. Elliptic shapes represent the footprint of implanted optical fibers. **f**, Reduced grooming during 5 Hz optogenetic stimulation of ChETA-expressing neurons in the mOFC (left). Quantification of reduced grooming (dots are individual animals) (right). **g**, mOFC photostimulation-induced reduction of grooming using 5, 15, and 120 Hz stimulation frequencies, where blocking the optic fibers abolished the effects on grooming. No effect of photostimulation on grooming was found in control animals (injected with stable fluorophore in mOFC), or after activating lOFC or M2. \**p*<0.05, #*p*<0.1.

## DISCUSSION

Although DBS is widely used for the treatment of otherwise therapy-resistant OCD patients, its mechanism of action remains poorly understood^41^. In particular, DBS-induced electrical artifacts have hampered the electrophysiological investigation of the brain during DBS^42,43^. Here, we overcame this limitation by using electrical-noise-resistant calcium imaging^28^ in SAPAP3^-/-^, a well-established mouse model for OCD^22^, to monitor cortex-wide population dynamics and single-cell activity in cortical and striatal regions, while systematically varying clinically relevant DBS parameters. We identified several “global” and “regional” effects of DBS. The DBS effects occurring uniformly across genotype and brain region, the “global” effects, were: Both direction and duration of recruitment of individual neurons were stable across different DBS-parameters and were dependent on network activity in the awake state; recruited neurons were scattered throughout brain regions and their numbers increased with DBS intensity; DBS recruited overlapping yet distinct neuron populations at therapeutic DBS intensities; and the relative balance between the number of neurons excited or inhibited by DBS (E/I balance) was generally preserved. The “regional” effects of DBS, those that varied depending on genotype and brain region, were: DBS modulated more mOFC neurons in SAPAP3^-/-^ (compared to WT); the E/I balance in mOFC leaned toward excitation; and DBS reduced the number of grooming-associated neurons specifically in mOFC of SAPAP3^-/-^. Furthermore, optogenetically mimicking DBS in the mOFC of SAPAP3^-/-^, but not lOFC nor M2, was sufficient to reduce compulsive grooming, providing insight into how DBS exerts its anti-compulsive effects.

Although DBS induced strong effects on brain and behavior, it did not alter basic brain function. For example, baseline activity (akin to resting-state activity) was not altered by DBS^37^. In addition, we found robust correlations between distance and activity of pairs of neurons (spatiotemporal correlations) in all recorded regions^38,39^, which were preserved during DBS. In contrast to unaltered basic brain function, compulsive grooming was reduced dose-dependently by alterations of current and pulse width, but not frequency, validating clinical practice of focusing on current and pulse width in search of optimal DBS parameters^9^. Clinical procedures employ chronically applied high-frequency continuous stimulation to treat patients, but recent work on DBS in Parkinsonian mice has suggested improved efficacy during cyclic (periodic DBS on and off) stimulation^44^. However, we found no effect of cyclic stimulation on compulsive-like behavior, which is consistent with a lack of therapeutic effect in a recent study in OCD patients^30^. In addition, we examined low-frequency stimulation, which also lacked neuronal and behavioral effects. Together, we conclude that chronically applied continuous high-frequency DBS effectively reduces compulsive-like behaviors without compromising basic brain function.

It has been suggested that IC-DBS exercises its efficacy via modulation of cortico-striato-thalamo-cortical circuits^2,12,45^, predominantly via recruitment of cortical regions^15^. Consistent with studies in patients^16–20^, we report IC-DBS neuronal recruitment in cortical and striatal regions in mice. We identified a number of global effects of DBS: 1) sustained neuronal recruitment was absent in anesthetized animals, illustrating that recruitment of sustained neurons is not simply a product of antidromic stimulation of IC white-matter, but instead depends substantially on network activity in cortico-striato-thalamo-cortical circuits present only in awake animals^36^. The small percentage of neurons that were nonetheless recruited in anesthetized mice (transiently excited neurons) did overlap with recruited neurons in awake mice, suggesting that transient neurons might be recruited by direct, antidromic stimulation of IC white-matter. 2) Neurons were recruited in a consistent fashion: the direction (excitation or inhibition) and duration (transient or sustained throughout DBS epoch) of modulation of individual neurons was consistent across different DBS intensities, suggesting that neurons have a predisposition to being recruited in a particular manner. 3) DBS recruited neurons did not cluster spatially but were randomly scattered throughout the imaged regions. 4) Similar to the intensity-dependent reduction in grooming, the number of neurons recruited was dose-dependent, suggesting the extent of neuronal recruitment relates to the behavioral effects. 5) Contrary to the *‘*inhibition hypothesis*’*^11^, which states that the mechanism of DBS is a functional lesion that results in local inhibition, we found both DBS-induced excitation and inhibition. Remarkably, in most regions, excitation and inhibition were relatively balanced (i.e., equal numbers of excited and inhibited neurons). Consistent with a recent study that reported DBS-evoked membrane depolarization that interfered with somatic action potentials^46^, this questions the DBS *‘*inhibition hypothesis*’*. 6) At therapeutic intensities (i.e., medium- and high-intensity DBS), only partially overlapping neuron populations were recruited. Taken together, we speculate that the therapeutic efficacy of IC-DBS in multiple psychiatric disorders (e.g., addiction, anorexia nervosa, and mood disorders)^2^ is associated with the aforementioned global effects of DBS. Thus, IC-DBS might modulate dysfunctional activity by exerting widespread effects recruiting neurons scattered throughout multiple frontal cortico-striatal circuits, without over-exciting neural tissue due to the maintained E/I balance. This rather non-specific recruitment of neurons and network nodes may reduce compulsivity as well as potentially alleviate symptoms associated with other psychiatric disorders.

The mOFC has been implicated in compulsive behavior in OCD^47,48^, and therapeutic IC-DBS alters mOFC activity^20^ and its projections^49^. Consistently, we identified a number of regional effects of DBS in SAPAP3^-/-^ that indicate an exceptional role for the mOFC in anti-compulsive effects of DBS: in comparison to all other recorded regions: 1) mOFC was predominantly recruited in SAPAP3^-/-^, 2) recruited mOFC neurons were more likely to be excited by DBS than inhibited, and 3) DBS reduced the number of grooming-associated neurons specifically in mOFC. Our optogenetic experiments demonstrate that excitation of mOFC neurons (mimicking DBS effects), but not lOFC nor M2 neurons, reduced excessive grooming (similarly to DBS). Previous rodent studies have linked subregions of the OFC to compulsive-like behaviors^24,50^. Ahmari et al report that repeated photostimulation of mOFC terminals in striatum in WT induced long-term enhanced grooming (but not during stimulation)^51^ and Burguière et al demonstrate that photostimulation of the lOFC and its striatal terminals reduced compulsive grooming in SAPAP3^-/-52.^ However, in contrast to our work, these studies applied optogenetic stimulation for extended durations (five and three min, respectively) and the target subdomains of the OFC were different to ours. These subdomains are functionally heterogeneous^53^, which may explain the modest discrepancy in results. In addition, a recent study reports ketamine-induced changes in dorsomedial prefrontal-cortex projections to the striatum in SAPAP3^-/-^ and direct photostimulation of this pathway resulted in reduced grooming^54^. This suggests that different therapeutic interventions and targets might normalize behavior via distinct prefrontal-cortex sub-circuits, where our data outlines a specific role for mOFC neurons in the therapeutic effects of DBS in compulsive behavior.

Our findings provide novel insights into the mechanism of action of DBS. We identified a large number of DBS effects which may be relevant for OCD as well as other psychiatric disorders treated with IC-DBS. These findings pave the way for further investigations into which effects are associated with what type of therapeutic utility. Our mOFC findings could inspire further clinical exploration of mOFC activity as a readout for DBS parameter optimization in OCD and other compulsivity disorders, and eventually may be used as a biomarker for closed-loop DBS. Such a biomarker has the potential to significantly shorten DBS-parameter optimization periods and improve DBS efficacy overall.

## Acknowledgements

We thank Dr. Tycho M. Hoogland, Andres de Groot, Mike Vink, and Joop Bos for technical support to perform imaging experiments. We are grateful to Ralph Hamelink, Dr. Nicole Yee, and Dr. Arthur S.C. França for technical assistance. We are thankful to Bart Kok and Makaela Weeda for assisting with histology. We thank Dr. Marcus H.C. Howlett, Dr. Matthijs G.P. Feenstra, and Dr. Ester Visser for their comments on the manuscript. We are grateful to the UCLA Miniscope team to develop and share the blueprints of the miniscopes. We thank the team of Dr. Karl Deisseroth for providing viral vectors. We acknowledge support from the Gravitation program of the Dutch Research Council grant, BRAINSCAPES (024.004.012), and a research grant from FFOR, the Foundation for OCD Research (I.W.).

## Author contributions

B.J.G.v.d.B designed the study, performed the experiments, curated, analyzed, and interpreted the data, wrote the manuscript, and finalized the paper. A.E.F. and P.A.R. assisted with experiments. E.H.v.B assisted with wide-field imaging experiments and edited the manuscript. A.P. assisted with analyses. D.D. edited the manuscript. I.W. designed and oversaw the study, interpreted the data, wrote the manuscript, and finalized the paper. All authors approved the work.

## Data availability

The data that support the findings reported in this article will be made available on Open Science Framework. The statistical analyses generated from the data are available on Open Science Framework (https://osf.io/w7qte/). Raw data are available from the corresponding author upon reasonable request.

## Code availability

The code used in this study will be made available on Open Science Framework.

## METHODS

### Experimental animals

Male and female SAPAP3 mutant mice (SAPAP3^-/-^, *n*=30) and their wild-type littermates (WT, *n*=28) were used for deep-brain stimulation (DBS) and imaging experiments, and Thy1-5.17 GCaMP6f mice (*n*=5)^31^ for DBS and wide-field experiments. For optogenetics experiments, male and female SAPAP3^-/-^ (*n*=22) were used. Animals were housed under a 12-hour reversed light/dark cycle with ad-libitum access to food and water (20-50 gr; 2-8 months with an average of 4 months). After surgery, mice were housed solitary and bedding material for nest building was provided. All experiments were in accordance with Dutch and European laws and approved by the Animal Experimentation Committee of the Royal Netherlands Academy of Arts and Sciences.

### Miniscope imaging and DBS surgery

For calcium imaging experiments employing miniaturized fluorescent microscopes (so-called miniscopes^35^), animals were anesthetized with isoflurane (3%), placed on an isothermal pad to maintain body temperature (37 °C), and placed into a stereotactic frame (Kopf Instruments, USA). Anesthesia was maintained at 1.5% isoflurane (flow rate: 0.6 ml/min O_2_/air mixture). The head was shaved and disinfected using 70% ethanol. The analgesic drug Metacam (10 mg/kg), nonsteroidal anti-inflammatory drug (NSAID) dexamethasone (diluted 2 mg/kg), and saline to prevent dehydration (100 ml/kg), were injected subcutaneously. An incision was made in the skin and lidocaine (100 mg/ml, Astra Zeneca, UK) was applied to the exposed skull and the periosteum removed. Skull was leveled in anteroposterior (AP) and mediolateral (ML) direction before marking the coordinates for unilateral GRIN-lens (left or right hemispheres were counterbalanced across animals) and bilateral internal-capsule (IC) DBS-electrode placements. GRIN lenses were placed in lateral (AP: 2.8 mm, ML: ±1.5 mm, DV: -2.2 mm) or medial (AP: 2.6 mm, ML: ±0.5 mm, DV: -2.2 mm) orbitofrontal cortex (lOFC: SAPAP3^-/-^ *n*=6, WT *n*=5; mOFC: SAPAP3^-/-^ *n*=5, WT *n*=5), prelimbic cortex (PL: SAPAP3^-/-^ *n*=5, WT *n*=5) (AP: 2.1 mm, ML: ±0.3 mm, DV: -1.9 mm), premotor cortex (M2: SAPAP3^-/-^ *n*=5, WT *n*=4) (AP: 2.3 mm, ML: ±0.35 mm, DV: -0.3 mm), dorsal (AP: 1.1 mm, ML: ±1.5 mm, DV: -2.7 mm), or ventral (AP: 1.1 mm, ML: ±1.1 mm, DV: -4.8 mm) striatum (DS: SAPAP3^-/-^ *n*=4, WT *n*=5, VS: SAPAP3^-/-^ *n*=5, WT *n*=4). After drilling holes, the skull was cleaned and dried, and covered with a layer of bone-attaching cement (SuperBond C&B, Sun Medical Co., LTD, Japan). To improve imaging quality^55,56^, we first slowly lowered a 25G needle (300 nm/min) using a custom-made stereotactic motorized arm (https://osf.io/w7qte/), left it positioned at target location for 5 min, and then slowly retracted it (300 nm/min). Next, we injected (200 nl/min) 500 nl virus (two injections of 250 nl) with a stereotact-mounted syringe (Hamilton, USA) ∼100 μm off imaging-target center and waited 5 min per injection to maximize diffusion of virus before retracting the syringe. For cortical-imaging experiments, we injected AAV-DJ-CaMKIIa.GCaMP6s (titre: 3×10^12 vg/ml, diluted 1:10, Stanford University Gene Vector and Virus Core) to express the calcium indicator GCaMP6s in pyramidal neurons^57^. For striatal-imaging experiments, we used AAV-DJ-hSyn-GCaMP6s (titre: 5×10^12 vg/ml, diluted 1:5, Stanford University Gene Vector and Virus Core) to target inhibitory neurons (of which 95% are medium spiny neurons)^57,58^. Using our motorized arm, we lowered (100 nm/min) the GRIN relay-lens (0.6 mm diameter, ∼7.3 mm long, Inscopix, USA), covered the gap between lens and skull with cyanoacrylate glue (Bison, The Netherlands), and used cranioplastic cement to secure the lens to the skull. Subsequently, we lowered custom-made DBS electrodes bilaterally into the IC (AP: -0.46 mm, ML: ±1.8 mm, DV: -4.6 mm)^27,59,60^, cemented DBS connectors to the skull using cranioplastic cement, and cemented a custom-made head bar to the skull. The GRIN relay-lens was covered and protected using Twinsil speed (Picodent GmbH, Germany). For M2 imaging, we additionally gave animals an subcutaneous injection of 15% D-Mannitol in saline (22 ml/kg) to aid diffusion of virus particles and reduce swelling of the brain^61^, injected four times 125 nl virus ∼100 μm off imaging-target center, and directly placed the GRIN objective-lens (1.8 mm diameter, Edmund Optics Ltd., UK) onto the brain. After surgery, animals received carprofen-analgesic containing drinking water (0.06 mg/ml) for three consecutive days. Animals were allowed to recover for one week.

### Wide-field imaging and DBS surgery

We used Thy1-5.17 GCaMP6f mice that express GCaMP6f throughout the cortex to image the entire dorsal cortex of one hemisphere^62,63^. Animals underwent similar surgery steps as described above (“Miniscope imaging and DBS surgery”). In addition, we applied a thin layer of cyanoacrylate glue (Bison, The Netherlands) to the skull, making the bone transparent^64^. One hole was drilled contralateral to the to be imaged hemisphere (AP: -0.7 mm, ML: -1.72 mm, DV: –5.48 mm) and the DBS electrode was inserted in a 40° angle to target the contralateral IC. For stability and to reduce light glare, we applied a layer of clear cement (SuperBond C&B, Sun Medical Co., LTD, Japan), followed by nail polish (Electron Microscopy Sciences, England). A head bar was placed posterior to lambda, and the outer edges of the clear skull were covered with a small wall of cement (Charisma, Kulzer, Germany) to prevent skin growth. After surgery, animals received carprofen-analgesic containing drinking water (0.06 mg/ml) for three consecutive days. Animals were allowed to recover for one week.

### Optogenetics surgery

For optogenetics experiments, SAPAP3^-/-^ underwent similar surgery steps as described above (“Miniscope imaging and DBS surgery”). In addition, we bilaterally injected (200 nl/min) 500 nl virus (two injections of 250 nl per hemisphere) with a stereotact-mounted syringe (Hamilton, USA) ∼100 μm off fiber-target center and waited 5 min per injection to increase diffusion of virus before retracting the syringe. We injected AAV-DJ-hEF1a.ChETA.eYFP (titre: 1.6×10^12 vg/ml, Stanford University Gene Vector and Virus Core) in the mOFC (*n*=7; AP: 2.6 mm, ML: ±0.5 mm, DV: -2.2 mm), lOFC (*n*=5; AP: 2.8 mm, ML: ±1.5 mm, DV: -2.2 mm), or M2 (*n*=5; AP: 2.3 mm, ML: ±0.7 mm, DV: -0.3 mm). To control for nonspecific light effects, we injected AAV8.CaMKIIα.mCherry (titre: 1×10^12 vg/ml, Zurich Viral Vector Facility) in the mOFC (*n*=5). We targeted custom-made optic fibers (FP200URT: 200 μm diameter, 0.5 NA, Thorlabs GmbH, Germany) 200 μm above the injection site (in a 10° angle for mOFC and M2), covered the gap between the fibers and skull with cyanoacrylate glue (Bison, The Netherlands), used cranioplastic cement to secure the fibers to the skull, and made the headcap light proof by painting the outside with black nail polish. After surgery, animals received carprofen-analgesic containing drinking water (0.06 mg/ml) for three consecutive days. Animals were allowed to recover for one week.

### Baseplating

Miniscopes were prepared for deep-brain imaging by drilling a hole in the housing, inserting a screw, and mounting a GRIN objective-lens with a custom-made 3D-printed spacer. Three weeks after surgery, animals were habituated to the experimenter for five consecutive days, followed by three days of habituation to the custom-made head-fixation device with running belt. Animals were head fixated to improve imaging quality during baseplating. During baseplating, the Twinsil-speed protective layer was removed and GRIN lens cleaned using lens paper. A baseplate was mounted onto the miniscope, which was mounted onto a stereotactic arm to hover over the implanted GRIN lens in order to find the best field-of-view. Once an optimal field-of-view was established (i.e., maximizing the number of visible neurons), the baseplate was cemented to the headcap and made light proof by painting the outside with black nail polish. The miniscope was removed and a protective cap installed on the baseplate to avoid damage to the GRIN lens.

### DBS application

DBS electrodes were custom-made and consisted of two bipolar twisted teflon-coated platinum/iridium wires (diameter: 112 μm; barewire: 75 μm; distance between the two poles: 0.5 mm). Mice were tethered to deliver DBS via a rotary joint (Adafruit, USA), allowing free unrestricted movement of the animals. DBS parameters were programmed in a digital stimulator (DS8000, WPI, USA) and generated by isolators (DLS100, WPI, USA). DBS settings were inspired by clinical parameters used in OCD patients at the Amsterdam University Medical Centers (Amsterdam UMC, location Amsterdam Medical Center, The Netherlands) and by our previous work^9,27^. DBS pulses were always biphasic, and depending on the experiment, one of the following three parameters was varied systematically while the other two were held constant: current (100, 200, or 300 μA), pulse-width (40, 80, or 160 μA), or frequency (60, 120, 180 Hz). The standard DBS parameters were (two of which were always held constant): 200 μA current, 80 μs pulse-width, and 120 Hz frequency. All mice underwent these three experiments, which occurred on different days (with weeks in between). In addition, we tried novel stimulation parameters: low frequency (1, 5, 20 Hz) and cyclic (DBS ON (200 μA, 80 μs, 120 Hz) for 10 s, OFF for 1, 5, or 10 s).

### Experimental setup for wide-field imaging experiments

Three weeks after wide-field imaging surgery, animals were habituated to the experimenter for five consecutive days, followed by three days of habituation to the custom-made head-fixation device with running belt. During imaging, mice were head fixated on a stable platform and placed under a wide-field fluorescence microscope (Axio Zoom.V15, ZEISS, Germany) to image the entire dorsal cortex of one hemisphere. Images were captured at 20 Hz (50 ms exposure), stored in 12-bit, 1600×1600 pixel images (∼15 μm per pixel), imaged by a high-speed sCMOS camera (pco.edge 5.5, PCO, Germany), and recorded using Encephalos software (Caenotec). Using an Arduino, the imaging computer triggered the digital stimulator to start and stop DBS.

### Experimental setup for DBS, miniscope imaging, and optogenetics experiments

Experiments were performed in two open fields (custom-made square, light-shielded Perspex boxes, 30 × 30 × 40 cm) housed inside sound-attenuated chambers. Videos were recorded with a Basler GigE camera (monochrome 1/2” Basler acA1300-60gm) attached to a Kowa lens (1/1.8”, F 1.6, 4.4–11 mm) and an IR-pass filter (43 mm, P = 0.75 mm), mounted in the center above (50 cm) the open field. Two infrared beams illuminated the open field from above and two infrared beams were mounted below the open field (IR-56, Microlight) to illuminate the open field through the transparent floor from below, creating strong contrast between animal and background. Behavioral videos were captured at 30 frames per s, with 1024×768 pixels, and stored in uncompressed AVI format using a custom-written script in the open-source software Bonsai^65^. A central computer controlled the cameras in both open fields (Dell T3500 workstation, Windows 7 64-bit), while also triggering the digital stimulator to start and stop DBS (via an Arduino), and triggered miniscope data acquisition cards to start and stop calcium imaging (via an Arduino), or triggered blue lasers (DPSS 473 nm, Shanghai Laser & Optics Century Co., Ltd., China) to start and stop photostimulation (via an Arduino). The central computer recorded the (behavioral) video frames and corresponding time stamps, sent TTL triggers (to trigger DBS, imaging, and photostimulation), and saved corresponding TTL trigger time stamps. Using these time stamps, we were able to align DBS, miniscope imaging, and photostimulation data to behavioral data.

### Miniscope-imaging sessions

Mice were habituated to the open fields (custom-made square, light-shielded Perspex boxes, 30 × 30 × 40 cm) for three sessions by placing them in the center of the open field and allowing them to move around freely for 30 min. Animals were head fixated briefly on the running belt for cleaning of the GRIN lens, attaching the miniscope to the baseplate, and connecting the animal to the DBS stimulator. We employed a 6-channel rotary joint (Adafruit, USA) to employ miniscope imaging and DBS in freely moving mice, which was held by a custom-made balancing arm to relieve weight of the animal*’*s head. To minimize bleaching of the calcium sensor in neurons, we imaged animals once a week, maxed out the miniscope sensor*’*s gain, and provided as little excitation LED as possible (0.5 - 10%). Each session consisted of four DBS blocks (e.g., during the current experiment: 0, 100, 200, or 300 μA stimulation conditions), with eight trials per block. Each trial consisted of 80 s of calcium imaging and 60 s of DBS, starting after 10 s of calcium imaging and ending 10 s before the end of calcium imaging. A fixed inter-trial-interval of 10 s was used between trials within a block, and an interval of 30 s between blocks. Systematic manipulation of a given DBS parameter (four blocks) was tested in a single recording session and animals were exposed to one recording session per week. The order of stimulation conditions within a session was determined by a Latin square design.

### Optogenetics sessions

Three weeks after surgery, animals were habituated to the experimenter for five consecutive days, followed by three days of habituation to the open fields (see above). Animals were tethered to a blue laser (DPSS 473 nm, Shanghai Laser & Optics Century Co., Ltd., China) via an optical rotary joint (1×2 fiber-optic rotary joint, Doric, Canada), which was held by a custom-made balancing arm to relieve weight of the animal*’*s head. Comparable to the DBS experiments, each session consisted of five optogenetics stimulation blocks (15 Hz for 5 s “transient” stimulation or 1, 5, 15, or 120 Hz for 60 s “sustained” stimulation), randomized across animals, with eight trials per condition. Each trial consisted of 120 s of (behavioral) video recording, starting 30 s before optical stimulation (which continued for 60 s in case of sustained stimulation and 5 s in case of transient stimulation) and lasted 90 s after the start of optical stimulation. We administered 5 mW of 473 nm blue light (10 ms pulse-duration) with different frequencies: either 5-s stimulation with 15 Hz to mimic transient activity, or 60-s stimulation with 1, 5, 15, or 120 Hz (4 ms pulse duration) to mimic sustained activity. To control for potential nonspecific effects of photostimulation on behavior, we 1) used animals injected with virus expressing a stable fluorophore (lacking an opsin) and 2) tested animals with ChETA in the mOFC in a condition where laser-light access into the brain was obstructed at the head cap (by a ferrule filled with black nail polish)^66,67^.

### Histology

Mice were deeply anesthetized using a lethal dose of pentobarbital, transcardially perfused with 4% PFA in PBS, and decapitated. Heads were submerged in 4% PFA for at least 24 h to preserve lens or fiber and electrode tracks. Subsequently, brains were removed, placed in 30% sucrose for cryoprotection, rapidly frozen using isopentane, and sliced on a cryostat (40 μm coronal sections, -20 °C). Coronal sections containing lens or fiber locations were stained with DAPI to visualize cell nuclei, mounted on glass slides, and imaged with an Axio Scan.Z1 slide scanner (ZEISS, Germany) to validate target location. Sections containing the IC were stained with cresyl violet and imaged with an Axioskop bright-field microscope (ZEISS, Germany) to validate DBS-electrode tip location. Headcaps (GRIN lens and head bar) were placed in acetone for 24 h, cleaned using acetone, ethanol, and lens paper, and reused.

### Modeled sphere of activation

We modeled the current spread around the tip of the DBS electrodes to validate stimulation of IC using the following formula:

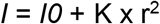

Where I is applied current (100 to 300 μA), I0 is amount of current needed to excite an axon (7-22 μA), given that the electrode touches the axon, K is a constant that describes how quickly the threshold current increases as the electrode is moved away from the axon (1292 μA/mm^2^), and r^2^ is squared distance between the axon and electrode. I0 and K are based on previous studies^68,69^. Using this formula, we found a sphere of activation with a diameter of 0.54 mm (low-intensity DBS) to 0.95 mm (high-intensity DBS).

### Grooming analyses

Grooming behavior was identified by a grooming classifier, as described previously^21^. We trained a Janelia Automatic Animal Behavior Annotator (JAABA) classifier to detect grooming in animals tethered to miniscopes. In short, animal behavior was video-taped and locomotion extracted using Bonsai^65^. Next, we extracted detailed frame-by-frame position information using the open-source software Mouse Tracker^70^, which was fed as input to the JAABA classifier^71^. A human expert observer trained the JAABA grooming classifier on 39.090 frames (19.133 grooming frames and 19.957 not-grooming frames) from eight short videos of SAPAP3^-/-^, which provide sufficient amounts of grooming frames. 1/7 Folding cross-validation showed that the classifier was able to reliably detect grooming with 82.3% sensitivity and with 74% specificity. To improve accuracy, we introduced a minimum bout length of 10 frames and set a higher threshold of 0.5 to detect grooming (to reduce false positives). Together, this resulted in 87.2% sensitivity and 92.1% specificity to detect grooming events (Supplementary Fig 1a,b,c). Grooming data were binned into 1 s bins and transformed into percentages. To examine whether DBS and photostimulation affected grooming, we used paired t-tests to compare grooming during DBS with grooming before DBS application. In order to explore the relationship between reduction in grooming and DBS electrode location, we calculated relative change in grooming [(grooming during DBS / grooming before DBS) -100] and correlated that with averaged electrode locations in both hemispheres [AP coordinates * DV coordinates].

### Wide-field calcium-imaging analyses

Images were binned into 800×800 pixels and converted to a 16-bit format. Images were spatially downsampled by a factor of 2 and registered to the first frame of that session or to the previous session. Subsequently, data were motion corrected within a session by first computing the 2D cross-correlation between the first frame and the remaining frames. Next, frames were rigidly shifted to achieve maximum correlation and inspected manually. Next, we smoothed the data using a Gaussian filter with a standard deviation of two pixels. Per pixel, we calculated relative DF/F, where DF is the activity at a given time and F is the mean activity per trial at 9 to 8.5 s before DBS application. Frames were aligned to the Allen Mouse Brain Common Coordinate Framework using Bregma, Lambda, and suture lines^33^. Regions were frontal cortex (FC), somatosensory cortex (SS), Visual cortex (VIS), and retrosplenial cortex (RSP). Pixels were averaged within these regions. Data were z-scored per region across the entire recording using the following formula:

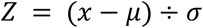

where *Z* is the standard score DF/F, *x* is the observed value, *μ* is the mean of the region, and *σ* is the standard deviation of the region. Sustained DBS-induced suppression was examined by statistically comparing z-scored DF/F signals during DBS (last 30 s of the DBS period) across DBS parameters. In addition, to compare between regions, we applied post-hoc tests corrected for multiple comparisons (Tukey*’*s HSD). Finally, we averaged signals across all regions to explore dose-dependent responses across the entire cortex.

### Miniscope calcium-imaging analyses: Preprocessing

Calcium-imaging videos were stored as uncompressed AVI files at a rate of 15 frames per s and binned per 1000 frames, and FIJI was used for raw-data inspection^72^. Per animal, we first concatenated all AVI files of a single session into a TIFF file and used logged timestamps of single frames to calculate missing frames (https://osf.io/w7qte/). Concatenated TIFF files were motion corrected using NoRMCorre^73^ and neuronal footprints and signals extracted using CNMF-E^74^. Data were spatially downsampled by a factor of 2 and we used the following CNMF-E parameters: gSig = 7, gSiz = 17, merge_thr = [1e-1, 0.85, 0], min_pnr = 7.4 (range 4 - 30), min_corr = 0.8 (range 0.8 - 0.95) (https://osf.io/w7qte/). After footprint and signal extraction, we manually cleaned the data using a custom-written user interface that showed spatial footprints of region-of-interests (ROIs) and temporal traces per ROI, calculated distance and correlation between ROIs, and provided the options to delete or merge ROIs (https://osf.io/w7qte/). ROIs with artificially small (∼half the size of average ROI) or large (∼twice the size of average ROI) spatial footprint were discarded as noise or background signal. ROIs with strong overlap (distance <15 pixels, correlation >0.8) were averaged and merged into a single ROIs. We generally used DF/F (C_raw), except for the isoflurane baseline-activity, and behavior-associated neuron analyses. For these analyses, we deconvolved each neuron*’*s DF/F using OASIS to get denoised traces, which were used to estimate calcium events^75^. Using the recorded time stamps, we aligned imaging data to DBS periods, as well as grooming periods. Data were z-scored per neuron across the entire recording session (see above for formula). All analyses were performed in Matlab (R2016b and R2020b, MathWorks Inc., USA).

### Miniscope calcium-imaging analyses: DBS-associated neurons

Trials were divided into pre-DBS baseline period (10 s) and three DBS periods (early (20 s), middle (20 s), late (20 s)). Neurons were classified as responders (recruited neurons) if signal during DBS significantly differed from the pre-DBS baseline signal (paired t-test across trials). For transient neurons, only the early DBS period differed from pre-DBS baseline. For sustained neurons, all three DBS periods differed from pre-DBS baseline.

### Miniscope calcium-imaging analyses: Consistency of modulation

Single-cell recruitment by DBS varied in duration (transient or sustained) and direction (excited or inhibited) of activity. To examine whether DBS would recruit neurons likewise across stimulation parameters (e.g., 100, 200, and 300 μA), we calculated the consistency of recruitment: We defined consistency of modulation as “neurons in one cluster” divided by “total neurons in that cluster”. “Neurons in one cluster” were all the neurons in a given functional cluster that do not fall into another functional cluster across DBS intensities, and “total neurons in that cluster” were all neurons across DBS intensities that were identified as such (e.g., the number of transient excited neurons not found to be sustained excited, or transient or sustained inhibited across other stimulation parameters divided by the total number of transient excited neurons found during DBS). We compared to chance using bootstrapping: From all recorded neurons, we randomly selected the number of neurons as found in the actual data, calculated consistency of modulation, repeated this 1000 times, and calculated summary statistics to compare to the true data.

### Miniscope calcium-imaging analyses: Overlap of neurons

We employed venn diagrams to express overlap of neuron populations. For two overlapping neuron populations, we employed conditional probability to assess the percentage of neurons recruited in condition B, given that they have been recruited in condition A (e.g., the percentage of neurons recruited by DBS under anesthesia, given they have been recruited by DBS in the awake state). Probability can be expressed using the following formula:

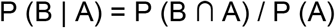

where P (B | A) are the neurons recruited in condition B, given that they have been recruited in condition A, P (B ∩ A) is the overlap of condition A and B, and P (A) are all neurons recruited in condition A. The overlap of neuron populations was compared to chance level using bootstrapping: From all DBS-recruited neurons across the DBS intensities, we randomly selected the number of neurons as found in each DBS intensity, calculated the overlap between intensities, repeated this 1000 times, and calculated summary statistics to compare to the true data.

### Miniscope calcium-imaging analyses: Regional baseline activity

To test whether DBS induced changes in regional baseline activity, we combined the cumulative density function (CDF) of calcium events and the frequency of activity into a single cell-activity index. We used the size of the calcium events (as used by CDF) but averaged across neurons and binned into one-minute bins (as used to calculate frequency of activity). Per animal, all single-cell deconvolved calcium events were summed up and averaged across neurons to compute the mean number of events per animal during the “no-DBS” block and the high-intensity DBS block. We did this for all animals to compare activity between the “no-DBS” block and the high-intensity DBS block.

### Miniscope calcium-imaging analyses: Clustering of recruited neurons

Miniscope imaging provides the spatial location of recorded neurons. To test whether DBS-recruited neurons would cluster (or maximally dispersed), we measured distance to the closest recruited neuron for each recruited neuron. Distance to the closest recruited neuron is important to avoid averaging out short (potential clusters) and long distances (potentially maximally dispersed), which would make the analysis unable to identify any spatial organization. To draw statistical conclusions about spatial modulation, we compared the true distance to chance level (bootstrap: from all recorded neurons, we randomly selected a subset equal to the number of DBS-recruited neurons, measured distance to the closest neuron, averaged over all neuron pairs, repeated this procedure 1000 times, and computed summary statistics to compare to the true closest distance). To validate this analysis method, we ran the analysis on simulated data and were able to identify different forms of clustering (single, multiple, and small clusters), a ring structure, maximally dispersed recruitment, and random recruitment.

### Miniscope calcium-imaging analyses: Behavior-associated neurons

Grooming, locomotion (extracted from Bonsai tracking data), and deconvolved data (calcium events, “S”) were binned into 333-ms bins (5 bins) to improve estimates of the neurons*’* activity^76^. Per session, we forced periods of grooming to have no locomotion (set value to 0 in the locomotion vector). Remaining locomotion values were split by their median: All values below median were considered stationary and all values above locomotion. Per neuron, we ran a Bayesian ANOVA with three categories of behavior: grooming, stationary, or locomotion. If the ANOVA was significant and Bayes factor > 3, we ran post-hoc tests, corrected for multiple comparisons (Tukey*’*s HSD), to compare activity during the three different behaviors. Classification of neurons was based on the following post-hoc comparisons:

1. grooming-associated neurons: grooming ≠ stationary & locomotion; stationary = locomotion;
2. locomotion-associated neurons: locomotion ≠ stationary & grooming; stationary = grooming;
3. grooming- and locomotion-associated neurons: grooming & locomotion ≠ stationary; grooming = locomotion;
4. not-associated neurons: not significant Bayesian ANOVA and Bayes factor < 1/3.

### Miniscope calcium-imaging analyses: Summary plots

We use summary plots to summarize main effects across different experiments. These plots present four dimensions: 1) The size of the “bubbles” depicts p-value (the bigger the bubble size, the lower the p-value), 2) color represents measured effect (e.g., change in cell-activity index), 3) columns generally represent the regions recorded (or optogenetic stimulation frequency), and 4) rows generally represent the different experiments (but other variables are possible, too).

### Statistical analyses

Data are presented as mean ± SEM. We used paired and independent *t-*tests, one- or two-way ANOVAs, and bootstrapping to determine statistical significance. A p-value of <0.05 was considered statistically significant. When appropriate, the alpha value was adjusted to correct for multiple comparisons (Holm-Bonferroni)^77^. For bootstrapping, we considered the true mean to be significantly different from a bootstrapped chance distribution if the 95th percentile ranges of the two distributions did not overlap. We computed p-values using:

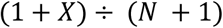

where *X* represents the number of overlapping data points between two distributions and *N*number of bootstraps^78^. We calculated effect size using Glass*’*s Delta:

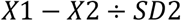

where *X*1 represents the population true mean, *X*2 bootstrapped distribution mean, and *SD*2standard deviation of the bootstrapped distribution. All statistical analyses were performed using Matlab (R2016b and R2020b, MathWorks Inc., USA).

