## Supplementary figures for "Unraveling the therapeutic mechanism of deep-brain stimulation"

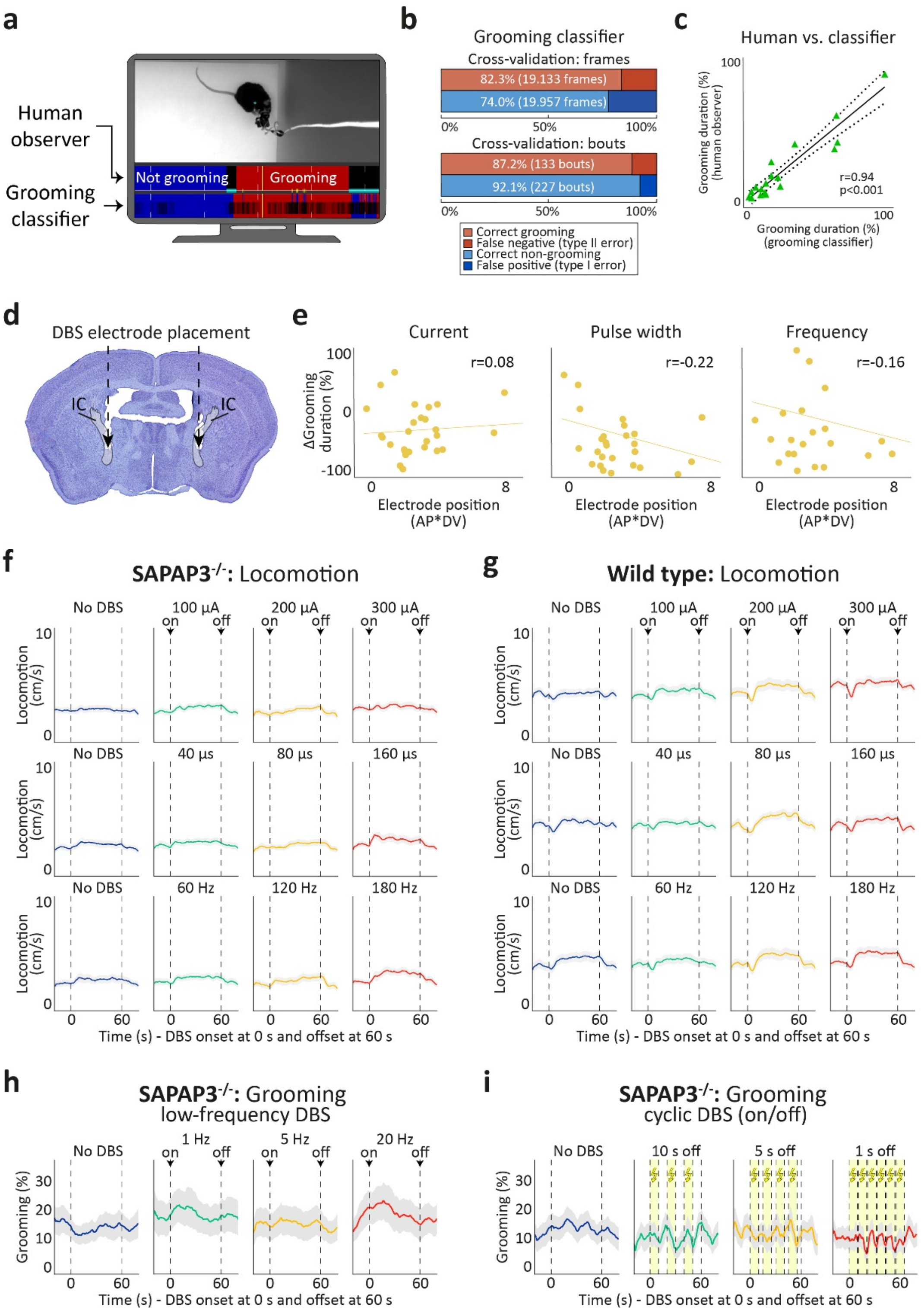


**Supplementary Fig. 1 | IC-DBS electrode location did not correlate with grooming reduction and did not affect locomotion. a**, Videos of behavior in the open field were used to train a grooming classifier using JAABA software[70]. Top row are behavioral assessments from a human observer, bottom row are predictions from the grooming classifier (red=grooming, blue=not grooming). **b**, Grooming classifier performance on single frames using k-fold cross-validation (true positive: 82.3%, true negative: 74.0%) (top). Grooming classifier performance on bouts (true positive: 87.2%, true negative: 92.1%) (bottom). **c**, Correlation of grooming scores from the human expert-observer versus the grooming classifier. **d**, Cresyl-violet staining with DBS electrodes in IC (gray) (representative example). **e**, IC-DBS electrode-position (two-dimensional anterior-posterior * dorsal-ventral) did not correlate with grooming reduction during high-intensity DBS. Dots are individual animals. **f**, No DBS-induced changes in locomotion in SAPAP3$^{-/-}$. **g**, No DBS-induced changes in locomotion in WT. **h**, Low-frequency IC-DBS stimulation (1, 5, or 20 Hz) did not affect grooming in SAPAP3$^{-/-}$. **i**, Cyclic IC-DBS stimulation (DBS on for 10 s and off for 10, 5, or 1 s) did not affect grooming in SAPAP3$^{-/-}$.

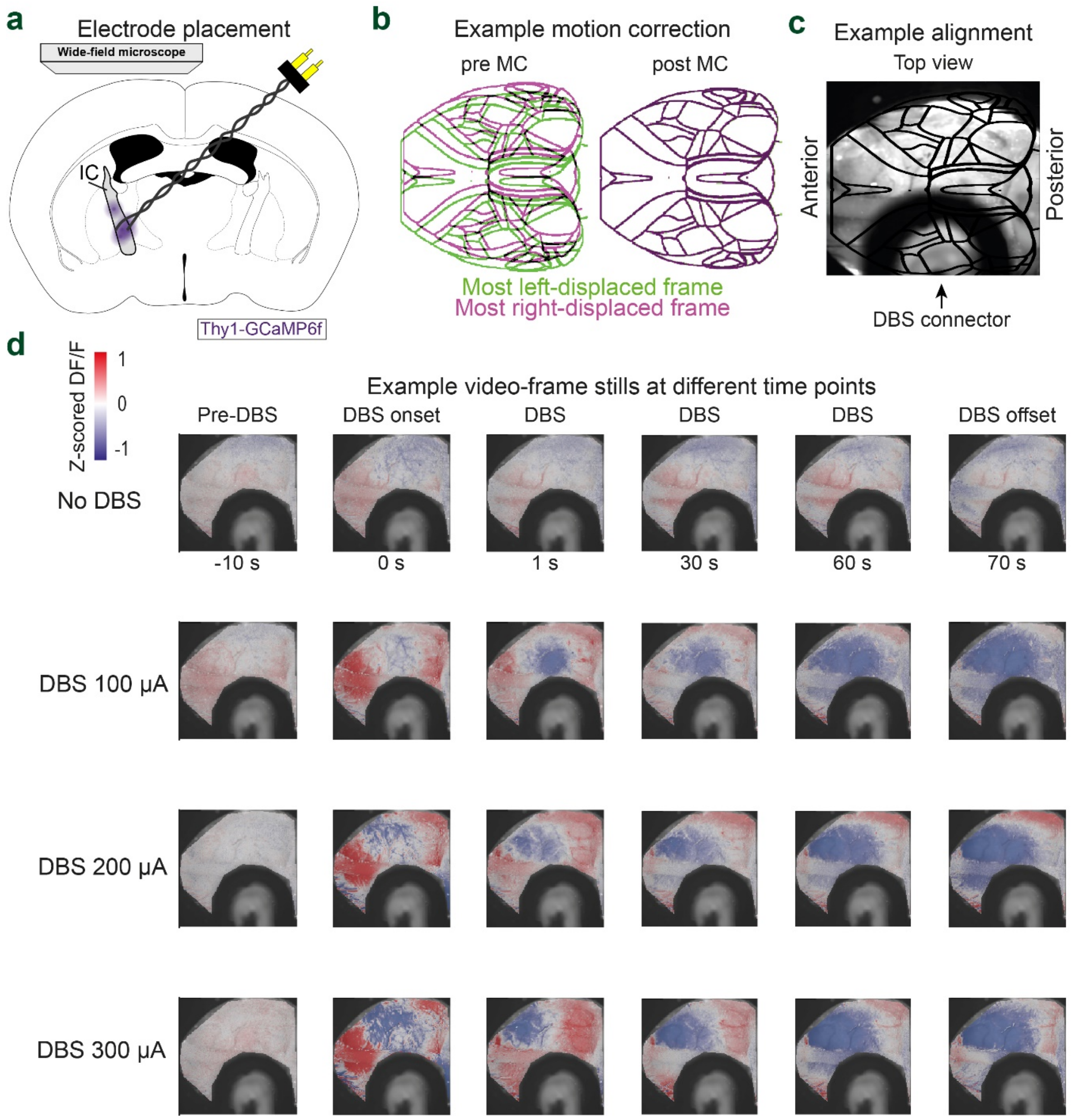


**Supplementary Fig. 2 | Representative data for wide-field imaging experiments. a**, Schematic[78] of the wide-field setup with DBS-electrode placement in IC (gray) of *Thy1*-GCaMP6f mice (*n*=5). **b**, Motion correction (MC) aligns individual wide-field video frames, as depicted by overlap of maximum deviating frames (left: green; right: pink) after motion correction. **c**, Alignment of a wide-field video frame to the Allen mouse-brain atlas (example video frame). **d**, Video frames of calcium dynamics during different DBS parameters (rows) across different epochs (columns).

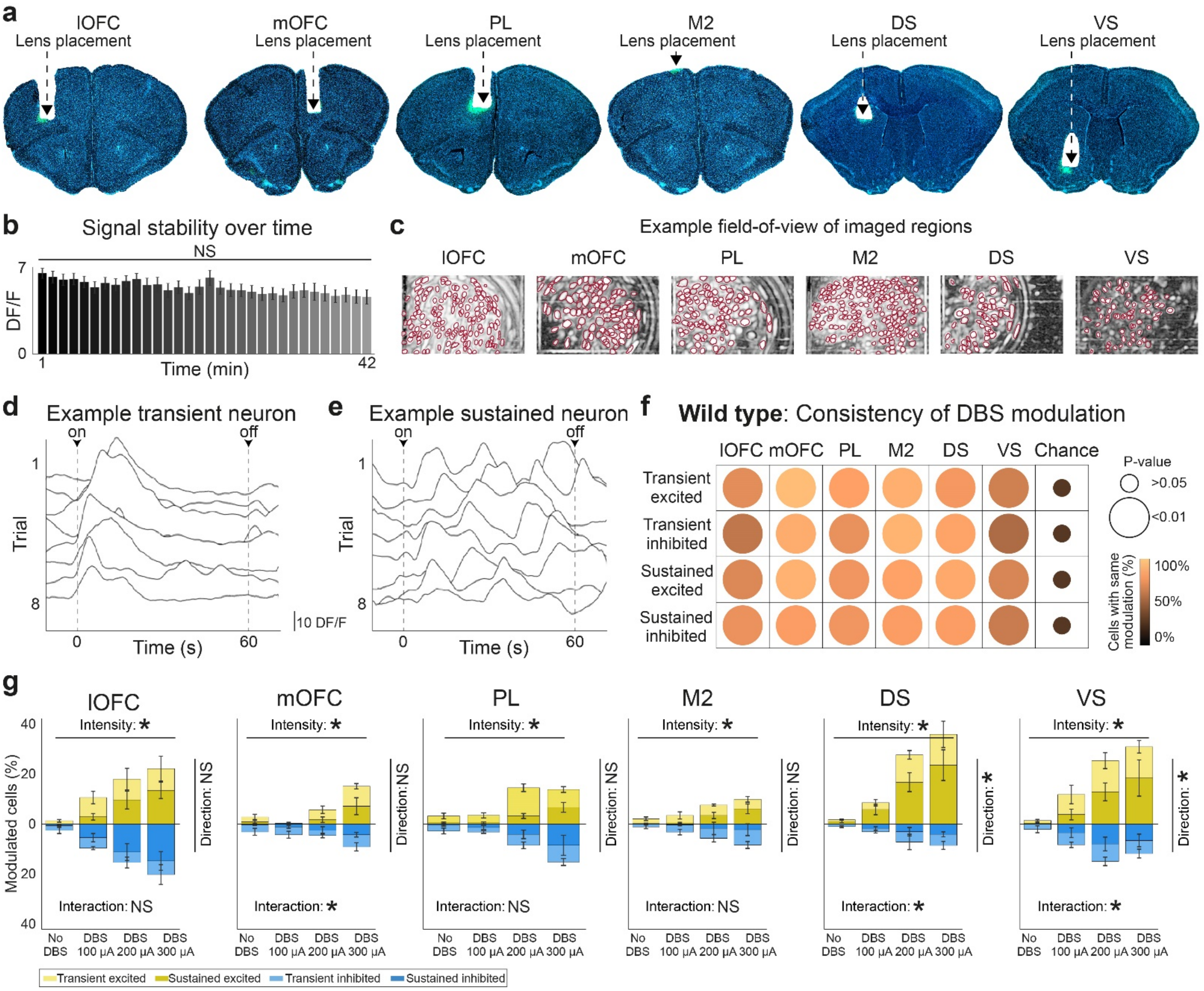


**Supplementary Fig. 3 | Modulation of cortical and striatal regions in WT by IC-DBS. a,** Histological verification of imaged regions (DAPI=blue, GCaMP6s=green). **b**, Miniscope calcium-imaging signal was stable across the entire recording session and did not exhibit bleaching. **c**, Maximum-intensity projection per region with overlay neuron outlines (in red). **d**, Example of a transiently excited neuron. For each trial, the neuron showed exclusively increased fluorescence upon DBS onset. **d,** Example of a sustainedly excited neuron. During each trial, the neuron showed calcium dynamics throughout DBS. **f**, Similar to SAPAP3$^{-/-}$ (Fig. 3g), consistency of modulation of DBS was significantly different from chance in all cortical and striatal regions recorded in WT. **g**, Similar to SAPAP3$^{-/-}$ (Fig 3f), we found dose-dependent recruitment of both excited and inhibited neurons by DBS (lOFC, PL, and M2). In other regions, we found differences in the number of neurons recruited by exciting or inhibiting their activity (mOFC, DS, and VS). *$p<0.05$, NS=not significant.

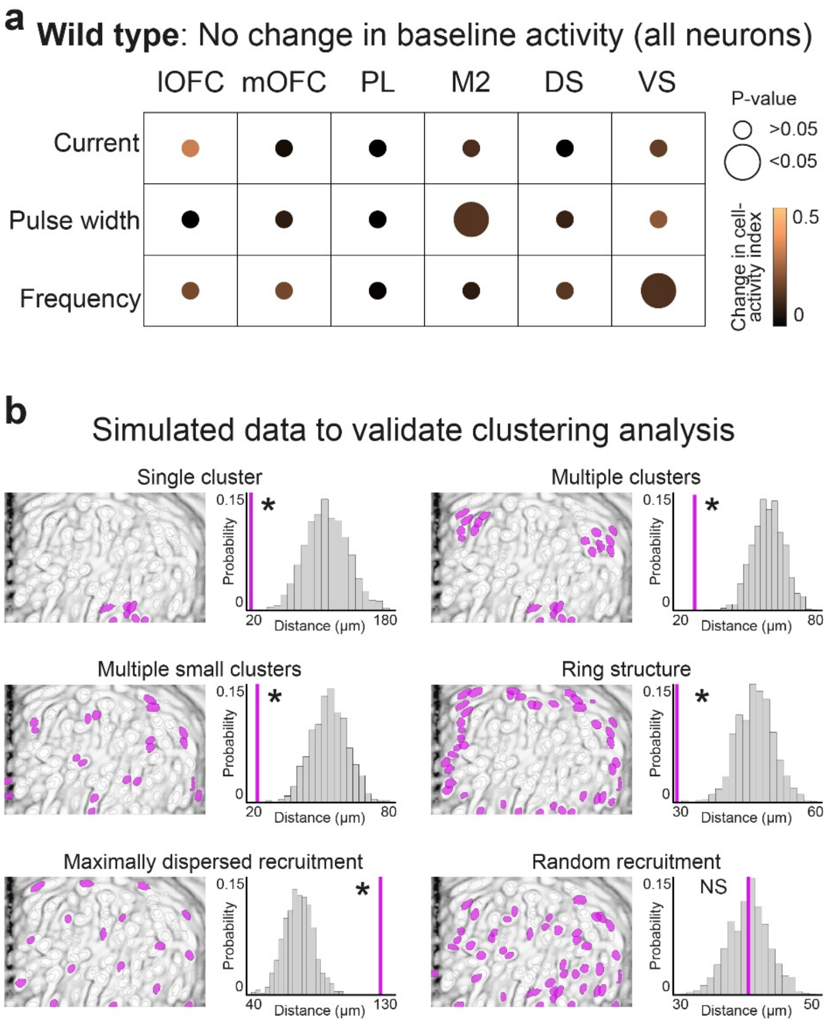


**Supplementary Fig. 4 | Baseline activity in WT and validation of cell-clustering analysis. a**, Similar to SAPAP3$^{-/-}$ (Fig. 4b), DBS did not alter cell-activity index in WT. **b**, Simulated data demonstrated detection of different forms of  clustering and dispersiveness. By comparing the distance to the closest neighbor to chance (bootstrap), the analysis can detect clustering as a single cluster, multiple clusters, multiple small clusters, or ring structure. In addition, dispersed recruitment (neurons distributed at equal distance throughout the field-of-view) and truly random recruitment could be detected. *$p$<0.05, NS=not significant.

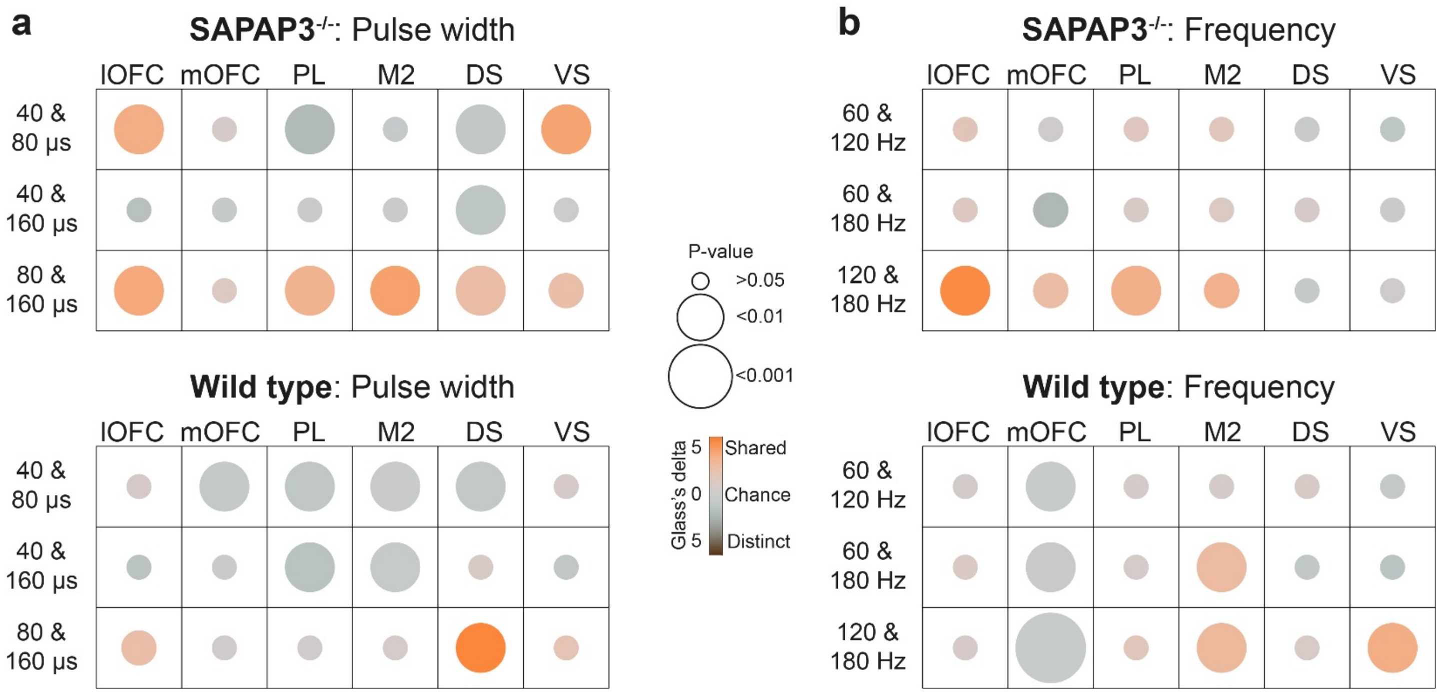


**Supplementary Fig. 5 | DBS recruits similar neuron populations during medium- and high-intensity DBS. a**, For the SAPAP3$^{-/-}$ pulse-width dose-response experiment, recruited neuron populations overlap significantly between 80 and 160 µs in almost all regions (top panel), but with little overlap in WT (bottom panel). **b**, Similar results as in a, but during the frequency dose-response experiment.

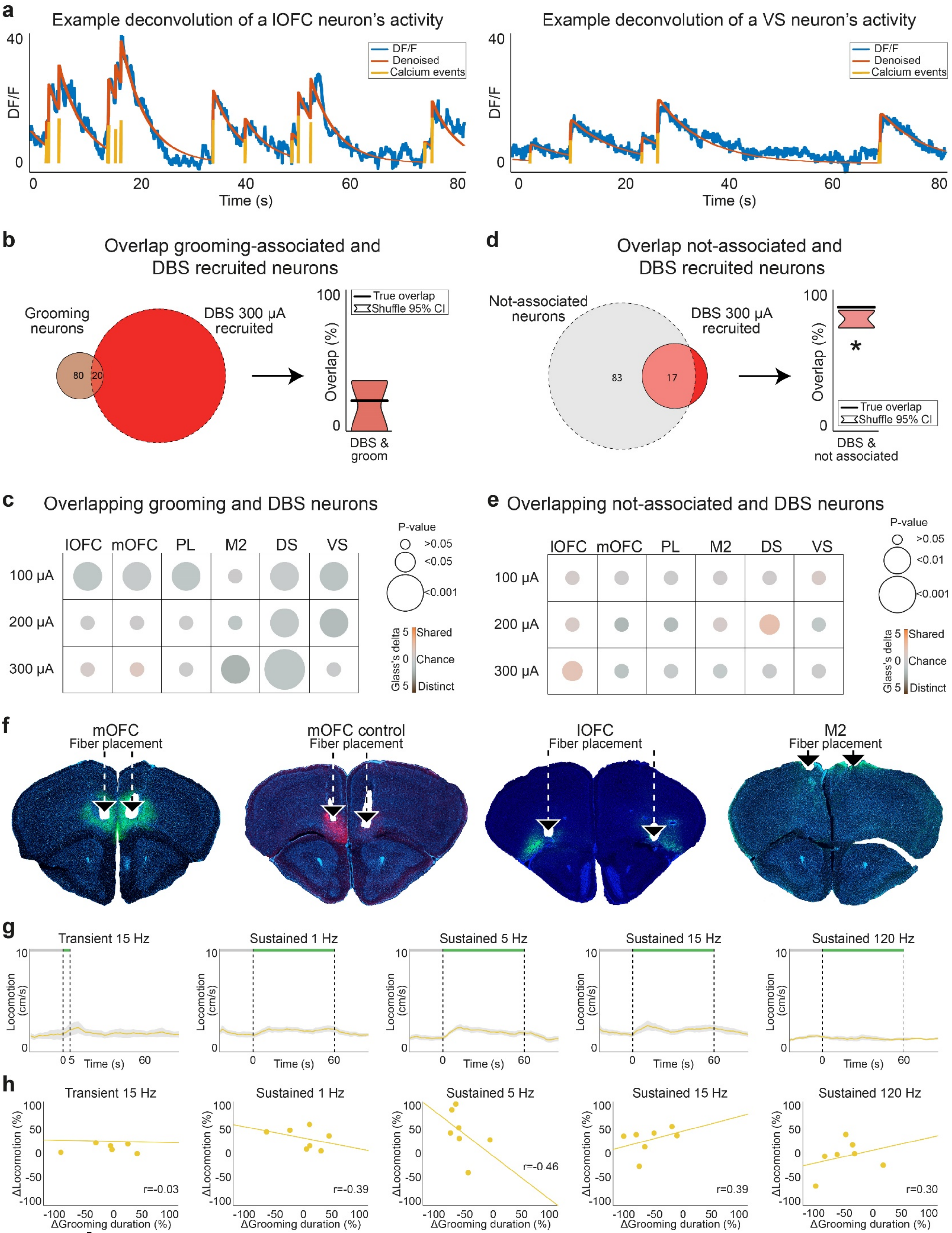
a
Example deconvolution of a lOFC neuron's activity
Example deconvolution of a VS neuron's activity
DF/F
Denoised
Calcium events
Time (s)
b
Overlap grooming-associated and DBS recruited neurons
Grooming neurons
DBS 300 μA recruited
True overlap
Shuffle 95% CI
Overlap (%)
DBS & groom
d
Overlap not-associated and DBS recruited neurons
Not-associated neurons
DBS 300 μA recruited
DBS & not associated
c
Overlapping grooming and DBS neurons
lOFC mOFC PL M2 DS VS
100 μA
200 μA
300 μA
P-value
Glass's delta
Shared
Chance
Distinct
e
Overlapping not-associated and DBS neurons
f
mOFC
Fiber placement
mOFC control
lOFC
M2
g
Transient 15 Hz
Sustained 1 Hz
Sustained 5 Hz
Sustained 15 Hz
Sustained 120 Hz
Locomotion (cm/s)
h
ΔLocomotion (%)
ΔGrooming duration (%)
r=-0.03
r=-0.39
r=-0.46
r=0.39
r=0.30

**Supplementary Fig. 6 | Non-specific recruitment of behavior-associated neurons by DBS and lack of locomotion effects during photostimulation.** **a**, Estimated calcium events (yellow) based on deconvolved trace (red) of fluorescent signal (blue) of a lOFC neuron (left) and a VS neuron (right). **b**, Overlap between grooming-associated neurons and DBS-recruited neurons (left) did not significantly differ from chance (bootstrap, right). **c**, DBS did not recruit grooming-associated neurons above chance (i.e., shared, overlapping neuron populations), or consistently below chance (i.e., distinct, unique neuron populations). **d**, Similar to panel b, but for not-associated neurons. **e**, Similar to panel c, but for not-associated neurons. **f,** Histological verification of stimulated regions (DAPI=blue, ChETA=green, mCherry=red). **g**, Locomotion during optogenetic stimulation of mOFC did not change across different stimulation frequencies. **h**, No significant correlation between reduced grooming and locomotion was found for any mOFC-photostimulation frequency. *$p<0.05$.
